# Nitrogen starvation response in hornworts and liverworts provides little evidence for complex priming to the cyanobiont

**DOI:** 10.1101/2024.05.22.595400

**Authors:** Yuling Yue, Gaurav Sablok, Anna Neubauer, Jaakko Hyvönen, Péter Szövényi

## Abstract

Mutualistic plant-microbe symbiotic interactions are thought to have evolved from a loose association between host plants and microbes when nutrients are limited. Therefore, the molecular network enabling intimate mutualistic plant-microbe symbioses may have evolved from a nutrient starvation response shared by all land plants. While the molecular link between nutrient status and symbiotic interaction is well-established, it remains poorly understood in some systems. This is especially true for the symbiotic associations between plants and cyanobacteria.

To test the conservation of the starvation network across land plants as well as to investigate the link between nutrient starvation and symbiosis initiation in the plant-cyanobacteria symbiosis, here we explore the transcriptional responses to nutrient starvation in two non-vascular plant species, a hornwort *Anthoceros agrestis* and a liverwort *Blasia pusilla*, forming plant-cyanobacteria endophytic symbioses. We observe a deep conservation of the systemic starvation response across land plants. However, very few if any components of the starvation network appear to be specific to cyanobacteria hosting plants, providing little evidence for extensive and specific priming to the cyanobiont. Moreover, we found that some bioactive molecules known to be important in initiating the plant-mycorrhiza and nodule-forming bacteria symbioses, may also have a similar role in plant-cyanobacteria symbioses.

**Highlight:** Our results suggest that the most critical step in establishing plant-cyanobacteria interactions using non-host plants is the attraction of the cyanobiont. This finding has significant impact on crop engineering.

## Introduction

It is hypothesized that the evolution of intimate mutualistic symbiotic interactions between plants and microbes occurred in multiple consecutive steps with the very first step entailing the capability of the host plant to attract microbes and initiate the symbiotic interaction in nutrient poor environments (Delaux and Schornack, 2021; Isidra-Arellano et al., 2021). Therefore, it has been proposed that the molecular network enabling and controlling intimate plant-microbe symbiotic interactions could have evolved from a general starvation response shared by all embryophytes (Isidra-Arellano *et al*., 2021; Shi *et al*., 2021, 2022). Indeed, it has been observed for decades that the establishment of plant-arbuscular mycorrhizal fungi (AMF) and plant-nodule forming bacteria symbiotic interactions can only be initiated under nutrient poor conditions. More specifically, increasing the concentration of phosphorus inhibits mycorrhizal infection of host plants (Balzergue et al., 2013). Similarly, nodule formation is triggered under nitrogen but also phosphorus starvation (Lu et al., 2020; Nguyen et al., 2021). This dependence is regulated by multiple molecular switches linking the genetic networks of the nutrient status of the host plant with those involved in the initiation and establishment of the symbiotic interaction (Shi et al., 2021; Jarratt-Barnham et al., 2022). For instance, plants able to form symbiotic interaction with AMF or nodule-forming bacteria release various compounds to the soil when kept under nutrient poor conditions. The best studied of these compounds are strigolactones and flavonoids (Misson et al., 2005; Hernández et al., 2007). Strigolactones are crucial for the initiation of plant-AMF symbioses by triggering the germination of fungal spores and growth of hyphae (Akiyama et al., 2005). Similarly, flavonoids released by the plant host are necessary to attract nodule forming bacteria by the plant host and to initiate the nodule-forming legume-bacteria interaction (Abdel-Lateif et al., 2012). Furthermore, master regulators of the phosphate/nitrogen starvation network have been shown to directly regulate many genes including members of the common symbiosis signaling pathway (CSP) required to prepare the host plant for accepting the symbiont including a specialized colonization interface and adjusted immune reaction (Shi et al., 2021). These observations are in line with the hypothesis that an abiotic nutrient starvation response is potentially conserved across embryophytes and might have been expanded to initiate and regulate plant-soil microbe interactions. Nevertheless, this hypothesis remains to be tested because information on the consequences of nutrient starvation is mainly available for vascular plants but missing for most non-vascular plant lineages.

While the molecular link between nutrient status of the host plant and establishment, as well as maintenance of the symbiotic interaction is well known in the plant-AMF and nodule forming bacteria symbioses, it is poorly understood in other plant-microbe symbiotic interactions. Some plants form endophytic, extracellular symbioses with cyanobacteria which are also under the control of the nutrient status of the host plant (Meeks, 1998; Xu and Wang, 2023; Álvarez et al., 2023b). On one hand, endophytic plant-cyanobacteria symbioses are similar to the plant-AMF and nodule forming bacteria symbioses because nitrogen starvation is necessary for the reconstitution of the symbiotic interaction in vitro as well as to maintain cyanobacterial nitrogen fixation (Enderlin and Meeks, 1983; Kimura and Nakano, 1990; CAMPBELL and MEEKS, 1992).

More specifically, upon nitrogen starvation the host plant is known to release the so-called hormogonia inducing factor (HIF), mobilizing and attracting the cyanobiont (Splitt and Risser, 2016; Nishizuka and Hashidoko, 2018; Hashidoko et al., 2019; Alvarenga et al., 2022). This response resembles that of the intracellular plant-AMF and -nodule forming bacteria symbioses in which flavonoids and strigolactones are secreted and sensed by the microbial partner. On the other hand, plant-endophytic cyanobacteria differ from the well-investigated plant-microbe symbioses (plant-AMF/-nodule-forming bacteria) in two important aspects. While most plant-endophytic cyanobacteria symbioses are extracellular and the cavities of the host plant containing the cyanobiont are present even in the absence of the cyanobiont (Meeks, 2003; Gorelova and Baulina, 2009; Xu and Wang, 2023), plant-AMF and -nodule forming bacteria symbioses are all intracellular and formation of host plant cellular structures essential for the colonization of the microbial partner must be induced by nutrient starvation and by contact with the microbial partner (Oldroyd, 2013; Ma and Chen, 2021). Therefore, nutrient starvation may induce a less complex genetic network in plant-cyanobacteria systems mainly related to the production of the HIF and immune system adjustment in contrast to intracellular symbioses (plant-AMF/-nodule-forming bacteria). Nevertheless, this hypothesis remains to be tested because the genetic network activated by starvation in plants forming endophytic symbiotic interaction with cyanobacteria is poorly characterized (Chatterjee et al., 2022).

To investigate these questions, we studied the transcriptional response to starvation in two non-vascular plant species representing ideal model systems of the plant-cyanobacteria symbioses using time-series RNA-seq data. We starved the hornwort model *Anthoceors agrestis* and the liverwort *Blasia pusilla*, with an estimated divergence time of 450 My representing two independently evolved realizations of the plant-cyanobacteria endophytic symbioses (Meeks, 1998; Duggan et al., 2013; Szövényi et al., 2015; Álvarez et al., 2023b). Then we compared gene expression changes associated with the starvation reaction in these two bryophyte species with data available for vascular plants to identify convergent and divergent transcriptomic responses (Fig. 1). We further used this data set to search for transcriptomic responses shared by the two cyanobacteria symbiosis forming organisms but missing from bryophytes and vascular plants unable to form symbiotic interaction, or only engaged in symbiosis with AMF and/or nodule-forming bacteria.

**Figure 1.**
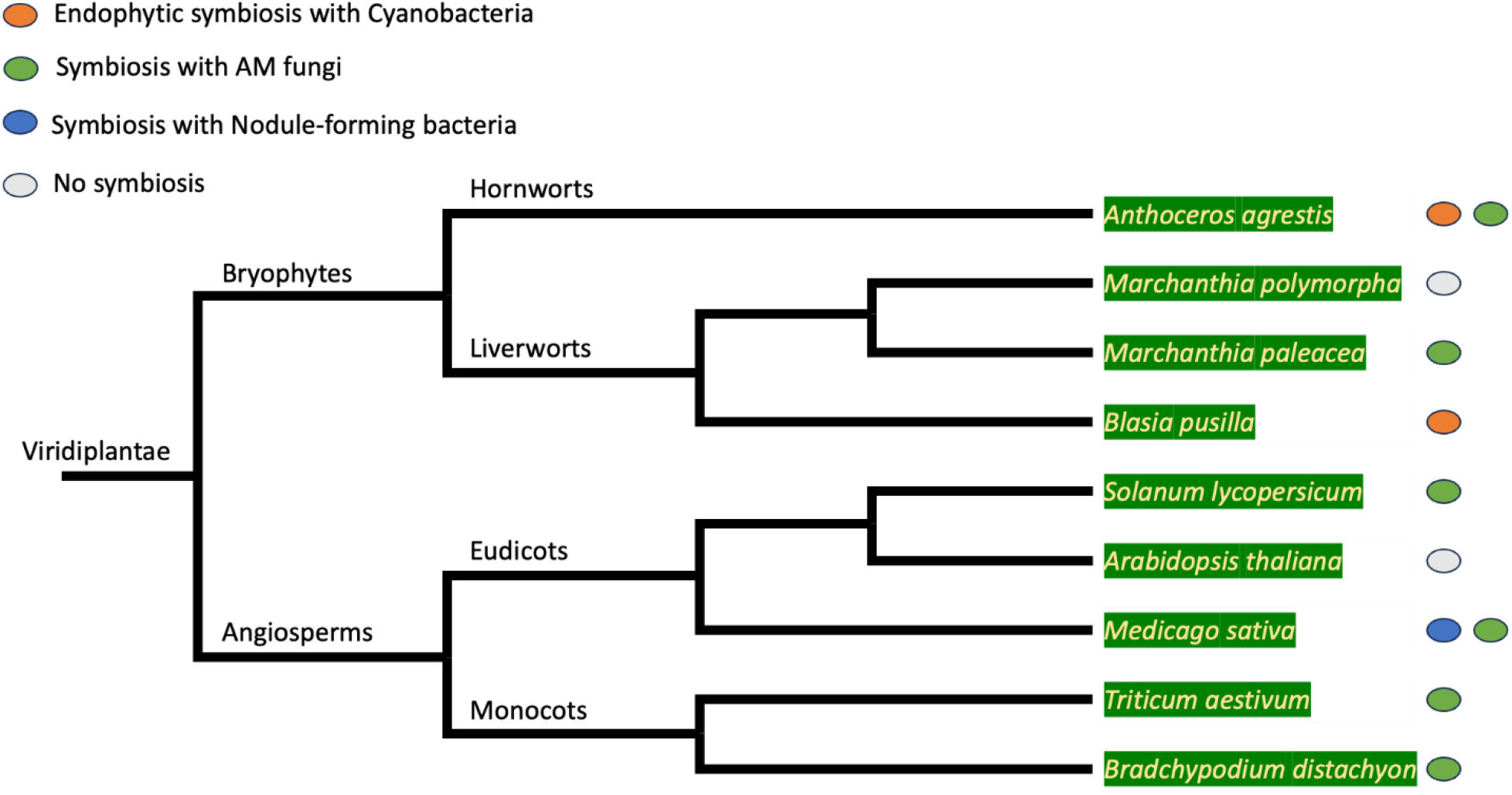
Phylogenetic relationship of the species used in this study and their symbiotic partners. Species were selected to represent four groups: species establishing endophytic symbiosis with cyanobacteria, species establishing symbiosis with AMF, species establishing symbiosis with nodule-forming bacteria, and species unable to form symbiosis with any of the three symbiotic partners. Ellipses after each species refer to the type of symbiotic interaction they are involved with.

## Materials and Methods

### Plant and cyanobiont material

We used the hornwort species *A. agrestis* and the liverwort model species *B. pusilla* to carry out the starvation experiment. The *A. agrestis* isolate was established from a single spore of the BONN isolate using culture conditions and techniques described earlier (Frangedakis et al., 2021). Young gametophytes were grown and vegetatively propagated for 1-2 months on solid KNOP (Frangedakis et al., 2021) medium in Petri dishes prior to initiating liquid cultures. The *B. pusilla* isolate was kindly provided by Elke Dittman (University of Potsdam). *B. pusilla* gametophytes were also precultured in Petri dishes on KNOP medium under the very same conditions described for *A. agrestis*. Prior to the starvation experiment we established liquid cultures for both species by dissecting Petri dish grown gametophytes into 0.5 mm fragments using forceps. We put gametophyte fragments into 200 ml liquid culture in a 500 ml flask containing BCD (Cove et al., 2009) and BG11 (Enderlin and Meeks, 1983) medium for *A. agrestis* and *B. pusilla*, respectively. To enhance plant growth, we kept the liquid cultures on an orbital shaker (150 rpm; SHKE2000-1CE; Thermo Fisher Scientific, Waltham, Massachusetts, USA) at 23°C under continuous light (17– 20 μmol·s^−1^·m^−2^, as measured using an LM2-1137 light meter with a Quantum Q37008 sensor [LI-COR Biosciences, Lincoln, Nebraska, USA]). Stock cultures of *A. agrestis* were subcultured in fresh media every two weeks by filtering the tissues using a cell strainer (BioSwisstec Premium Cell Strainer, 100 µm pore size). As cyanobiont, we used *Nostoc punctiforme* ATCC 29133. We precultured *N. punctiforme* on solid BG11 medium in Petri dishes for 2-3 weeks and inoculated 50 ml BG11 in a 125 ml flask. These liquid cultures were kept under the same conditions as the hornwort and the liverwort cultures.

### Starvation experiment

To start the starvation experiment, we collected two cell strainers (BioSwisstec Premium Cell Strainer, 100 µm pore size) of plant tissue for each plant species (corresponding to a ∼169 mg dry weight) and transferred them into one 500 ml Erlenmeyer flask, in total we used 12 flasks. Six out of the 12 flasks contained 200 ml fresh media without combined nitrogen (BCD-N/BG11-N, “starvation flasks”, details are provided in the Table S1), while the other six flasks were filled with 200 ml fresh media with combined nitrogen (BCD/BG11, “control flasks”, details are provided in the Table S1). To monitor plant response to starvation, we sampled thallus tissues from the starvation flasks after 1,2,3,7,8,9, and 10 days and from the control flasks after 1,2,3,7, and 10 days of inoculation (details see in Table S1). On each date, we collected ∼110 mg fresh weight of plant tissue (corresponding to ∼3.5 mg dry weight) in five biological replicates for both species.

### RNA extraction and sequencing

We immediately submerged collected thallus tissues of *A. agrestis* into RNA*later*^®^ (Sigma-Aldrich, product number: R0901-100ML) and stored them at 4°C overnight. On the next day, we removed the tissues from the RNA*later* solution, flash-froze them in liquid nitrogen, and stored them until RNA extraction at −20°C or −80°C in the freezer. For *B. pusilla*, we directly flash-freeze collected thallus tissues in liquid nitrogen. To extract RNA, we grinded the plant tissue into a fine powder with a pestle and a mortar in the presence of liquid nitrogen and processed the samples using the Spectrum^™^ Plant Total RNA-Kit from Sigma-Aldrich (product number: STRN250), including on-column DNA digestion using the on-column DNAse kit (Sigma-Aldrich, product number: DNASE70-1SET). The Agilent 2100 bioanalyzer (Agilent, Santa Clara, CA) and nanodrop were used to test the integrity, purity, and concentration of the RNA samples. For each sampling point we kept the three biological replicates with the best RNA qualities. The 42 mRNA libraries for *B. pusilla* and 30 mRNA libraries for *A. agrestis* were prepared using NEBNext® Ultra Directional RNA Library Prep Kit and paired end sequencing was performed on Illumina NovaSeq6000 (2 × 150 bp) by Novogene UK, Cambridge, London.

### Draft assembly of the Blasia pusilla genome

In-vitro grown gametophytic cultures of *B. pusilla* on MS medium (Murashige & Skoog Medium (MS1), 2008) were used for genomic DNA extraction using the plant-EZ DNA extraction kit and were sequenced using PacBio-SMRT at the Institute of Biotechnology, University of Helsinki, Finland. Reads mapped (using BLASR (Chaisson and Tesler, 2012)) to organelle and cyanobacterial genomes were removed. In addition to the PacBio sequencing, short read sequencing was done to support hybrid genome assembly and polishing. Short reads were trimmed using Fastp (Chen et al., 2018) and assembled together with the filtered PacBio reads using the hybrid genome assembler Masurca (Zimin et al., 2013). The hybrid genome assembly was polished with cleaned short reads using Pilon (Walker et al., 2014) with a minimum read depth of 10. Finally, we used blobtools v.1.1.1 (Laetsch and Blaxter, 2017), the NCBI nr database and the average coverage of Illumina reads for each scaffold to remove scaffolds of contaminant origin. After visual assessment we kept scaffolds with a taxonomic assignment of viridiplantae or streptophyte. Scaffolds with other taxonomic assignments were discarded. Completeness and redundancy of our assembled genome was assessed by searching translated peptide sequences of the reconstructed gene set against the hidden Markov model of Benchmarking Universal Single-Copy Orthologs BUSCO V.5 (Seppey et al., 2019) using the Viridiplantae and Embryophyta datasets retrieved from OrthoDB v11 (Kuznetsov et al., 2023).

### Creating a comprehensive transcript dataset for Blasia pusilla

We combined all RNA-seq data and the draft genome sequence to create reference transcripts for *B. pusilla* which was used later for gene expression estimation. We generated a genome-guided as well as a *de novo* TRINITY transcriptome assembly using all collected RNA-seq reads (42 RNA libraries) (Grabherr et al., 2011). We used the PASA pipeline (Haas et al., 2008) to combine the *de novo* and genome-guided assemblies into a non-redundant set of transcripts and putative genes. Using this approach, we created a transcript to gene mapping file and a non-redundant set of transcripts. We ran all transcripts through Transdecoder (https://github.com/TransDecoder/TransDecoder) to obtain their best ORF and peptide translation. To reduce the number of potential transcripts and gene models, we discarded all putative gene ids without at least one complete ORF prediction. To identify potential contaminants in this filtered transcript file, we selected the longest ORF for each putative gene and searched them against the Eggnog database in TRAPID 2.0 (Bucchini et al., 2021) and assessed their taxonomic assignment. The initial run indicated that most of the contamination concerned yeast. To remove these transcripts, we searched all transcripts against the full transcriptome of *Saccharomyces cerevisisae* S288C (assembly R64) transcripts using blastn version 2.12.0+ (Altschul et al., 1990) and discarded all transcripts passing the following filtering criterium: -evalue ≤ 0.0001, similarity value ≥ 90%, and query coverage ≥ 80%. (Yeast transcriptomics data was downloaded from here: https://ftp.ncbi.nlm.nih.gov/genomes/all/GCF/000/146/045/GCF_000146045.2_R64/GCF_000146045.2_R64_rna.fna.gz) After that, we ran a further taxonomic assignment using TRAPID 2.0 and removed another smaller set of genes mapping to Viruses, Bacteria, Archeae, Opisthokonta, and other Eukaryota. The final assembly had 22635 genes represented by 156239 transcripts.

The quality of the assembled transcripts was evaluated using three methods: (1) by assessing the read content of the transcriptome; (2) by calculating the ‘gene’ Contig Nx Statistic; and (3) the completeness of the transcript dataset. To assess the read content of the transcript assembly, we counted the number of overall aligned read pairs to our assembled transcripts using Bowtie2 (Langmead and Salzberg, 2012). We computed the conventional Nx length statistic using the Trinity toolkit utilities (TrinityStats.pl). Finally, we assessed the completeness of the transcript dataset by BUSCO V.5 (Seppey et al., 2019) using both the Viridiplantae and Embryophyta references retrieved from OrthoDB v11 (Kuznetsov et al., 2023).

### Expression quantification and time-series analysis

We applied the transcript quantification method in the mapping-based mode implemented in Salmon 1.4.0 (Patro et al., 2017), the other parameters were set to default. As reference we used the *B. pusilla* transcriptome set (assembled in this study), and the *A. agrestis* BONN cDNA sequences (Li et al., 2020). After running Salmon, we imported the quantification files into R with the tximport package (Soneson et al., 2015). The median of ratios method of normalization was used to normalize expression estimates. Prior to further statistical analyses, we removed transcripts from the data set for which the sum of the normalized expression counts over all samples was ˂ 10. We summarized gene expression values at the gene level for both species.

To gain insight into the overall qualitative change of gene expression associated with starvation in *B. pusilla* and *A. agrestis*, we carried out principal component analysis (PCA) (Brand, 2013) on normalized and standardized RNA-seq expression data. To identify genes significantly responding to nitrogen starvation in *A. agrestis* and *B. pusilla* plants, we employed the approach provided in maSigPro version 1.66.0 (Nueda et al., 2014), an algorithm modeling gene expression time series data with GLMs (general linear models). We chose the backward selection algorithm with an alpha value of 0.05, a Q-value threshold of 0.05 and a polynomial degree of 3. The expression data comprised RNAseq data of seven and five time points for *B. pusilla* and *A*. a*grestis* respectively. We clustered the significant (Q ≤ 0.05) DEGs into different profiles using the “hclust” cluster method in maSigPro and the Mclust option estimating the optimal k, choosing k.mclust=TRUE. Finally, we have tested different numbers of k to obtain a robust clustering of expression profiles by comparing the clusters at each step of k with the previous ones using the command “see.genes (get$sig.genes, k=…).”.

### Comparative transcriptomic analysis

To compare the gene set reacting for nitrogen starvation in *A. agrestis* and *B. pusilla* and that of vascular plants, we created orthogroups using Orthofinder v2.5.5 (Emms & Kelly, 2019) with default options. We included symbiotic and and non-symbiotic bryophyte and vascular plant species covering a range of taxonomic diversity increasing the accuracy of the orthofinder algorithm (full species list is provided in Table S2). Furthermore, we collected transcriptomic data in response to starvation for bryophytes and vascular plants able or unable to establish symbiotic interaction with AM fungi, nodule-forming bacteria, cyanobacteria, or a combination of these partners. More specifically, we obtained data for non-symbiotic species (*Marchantia polymorpha*, *Arabidopsis thaliana)*, species involved in interaction with AMF (*Marchantia paleacea, Triticum aestivum, Solanum lycopersicon, Brachypodium distachyon, Medicago sativa*) or nodule-forming bacteria (*Medicago sativa*). We retrieved the up and downregulated gene lists for all these species but *M. polymorpha* from the supplementary material of the original publications (see Table S3). For *M. polymorpha* we used DESeq2 (Love et al., 2014) method to test for differential gene expression (details are provided in the Table S4). Finally, expression information was added to the orthofinder results and orthogroups with convergent or divergent expression responses were identified. For *A. agrestis* and *B. pusilla*, up and downregulated genes were identified in the maSigPro analyses explained above. For the other species *M. paleacea, A. thaliana, B. distachyon* we retrieved the up and downregulated set of genes as they were given in the corresponding publications. For species such as *T. aestivum*, *M. sativa* and *S. lycopersium* in which transcriptomic response was recorded in multiple tissue types, we did the following: we selected the most significant differential expressed genes among all the tissues tested. If data for different tissues were available and the reaction of genes was tissue-dependent, we selected the greatest absolute log2 fold change value.

### Functional enrichment

To functionally identify and characterize the broad biological pathways and processes involved in nitrogen starvation, we obtained GO and KEGG annotations for the *A. agrestis* and *B. pusilla* gene set using Trapid 2.0 (see Table S5-S8). We carried out gene ontology and KEGG enrichment analysis of up and downregulated genes using the Parent-Child Algorithm and a Fisher’s exact test implemented in the TopGO package 2.46.0 (Alexa, 2006). We also carried out manual annotation of genes showing differential expression using the Plaza 5.0 database (https://bioinformatics.psb.ugent.be/plaza/). We retrieved transcription factor annotations and genes involved in hormone synthesis from the hornwort genome paper (Li et al., 2020). Furthermore, we also processed the full proteome of both species using the PlantTFBD website (https://planttfdb.gao-lab.org/)(Tian et al., 2020) to assign genes to transcription factor (TF) families (see Table S9).

## Results

### Draft genome assembly of Blasia pusilla

To aid our analyses, we generated a draft genome assembly for the *B. pusilla* with a combination of short and long-read data (see Methods section). After removing potential contaminants with Blobtools, our assembly consisted of 1456 scaffolds/contigs with a total length of ca. 347 Mbp (see details in the Table S10 and the final genome assembly on Figshare) which fits to the genome size estimated by previous c-value analyses (Renzaglia et al., 1995). Our results showed that the blobtools filtered draft genome assembly was of good quality containing 71%/79.7% (Embryophyta/Viridiplantae dataset) of the BUSCO genes. These BUSCO values are close to the figures obtained for high-quality genome assemblies generated using long-read technologies (*Marchanthia polymorpha*, *Physcomitrella patens*, *Pleurozium schreberi,* and *Funaria hygrometrica*) (Bowman et al., 2017; Lang et al., 2018; Pederson et al., 2019; Kirbis et al., 2020). Furthermore, over 96% of our RNA-seq reads could be mapped to the draft genome implying that the coding part of the genome is well covered (<u>Supplementary File 1</u>).

### Comprehensive transcriptomic data set

Initial annotation of the draft genome using braker1 and RNA-seq evidence resulted in a considerable proportion of fragmented gene models. Furthermore, we observed a large number of highly expressed gene models that were present in the transcriptome assemblies but missing from the genome-based braker annotation. Therefore, we decided to carry out a genome-guided as well as a *de novo* transcriptome assembly and combined their results. After contaminant filtering (eg. yeast Viruses, Bacteria, Archeae, Opisthokonta, and other Eukaryota) and removing gene models lacking at least one complete CDS sequence, the final assembly consisted of 22636 genes that were represented by 156239 transcripts. Completeness of the contamination filtered gene set (longest ORF per gene) was superior to that obtained by conventional annotation of the genome (see values above 71% and 79.7%) and captured 85.1% and 97.7% of the Embryophyta and Viridiplantae BUSCO gene set, respectively (The final transcriptome assembly is available on Figshare). The transcriptome assembly was of high quality based on all three statistics assessed (<u>Supplementary File 2</u>).

### Nitrogen starvation induces transcriptional reprogramming in both bryophytes investigated

Visualization of gene expression changes under nitrogen starvation over the time course of the experiment (standardized and centralized PCA) indicated considerable transcriptional reprogramming in both species (Fig. 2). More specifically, control and starvation samples formed two well distinguishable groups in both species after seven days. While transcriptional reprogramming occurred in both species, the temporal reactions to starvation differed considerably. In the hornwort (*A. agrestis*) the difference between control and starvation samples became considerable for the last two sampling points of the experiment. By contrast, in the liverwort (*B. pusilla*) global transcriptional changes were already detectible after a single day of nitrogen starvation. Quantitatively, we found that 20737 *A. agrestis* gene models were expressed in at least one of the nitrogen starvation samples, which is about 80.5% of the 25837 currently annotated gene set (Li et al., 2020) (Table S11, S12). Similarly, of the total 22636 gene models 22331 (98.6%) showed expression during nitrogen starvation in *B. pusilla* (Table S13, S14). Our statistical analysis indicated (MaSiPro Q≤0.05) that a similar number of genes, 1522 (7.34% of the expressed genes) and 1023 (4.58fig. 1% of the expressed genes), reacted to nitrogen starvation in *A. agrestis* and *B. pusilla*, respectively (Fig. 3 and Fig. 4).

**Figure 2.**
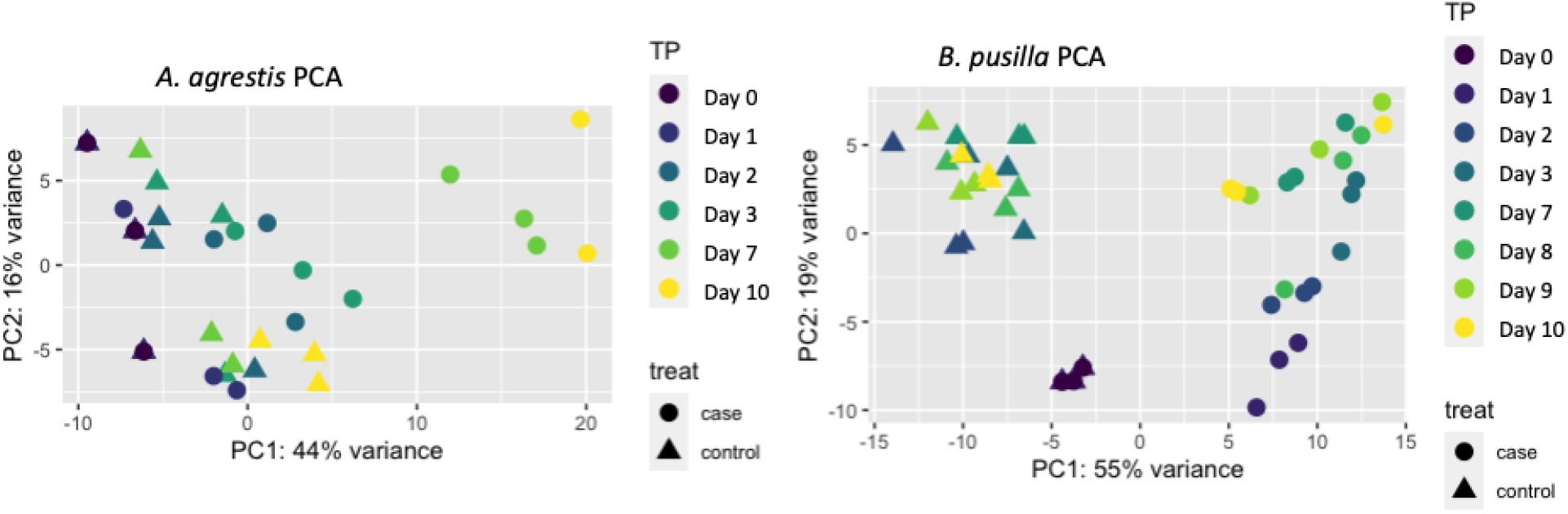
Principal component analysis (PCA) of overall transcriptomic response to nitrogen-starvation in *A. agrestis* (36 samples) and B. *pusilla* (48 samples). Each data point (triangle/dot) refers to a sample. Case (solid dot): samples starved of combined nitrogen; Control (solid triangle): grown in full media (see materials and methods); TP: sampling days after starvation.

**Figure 3.**
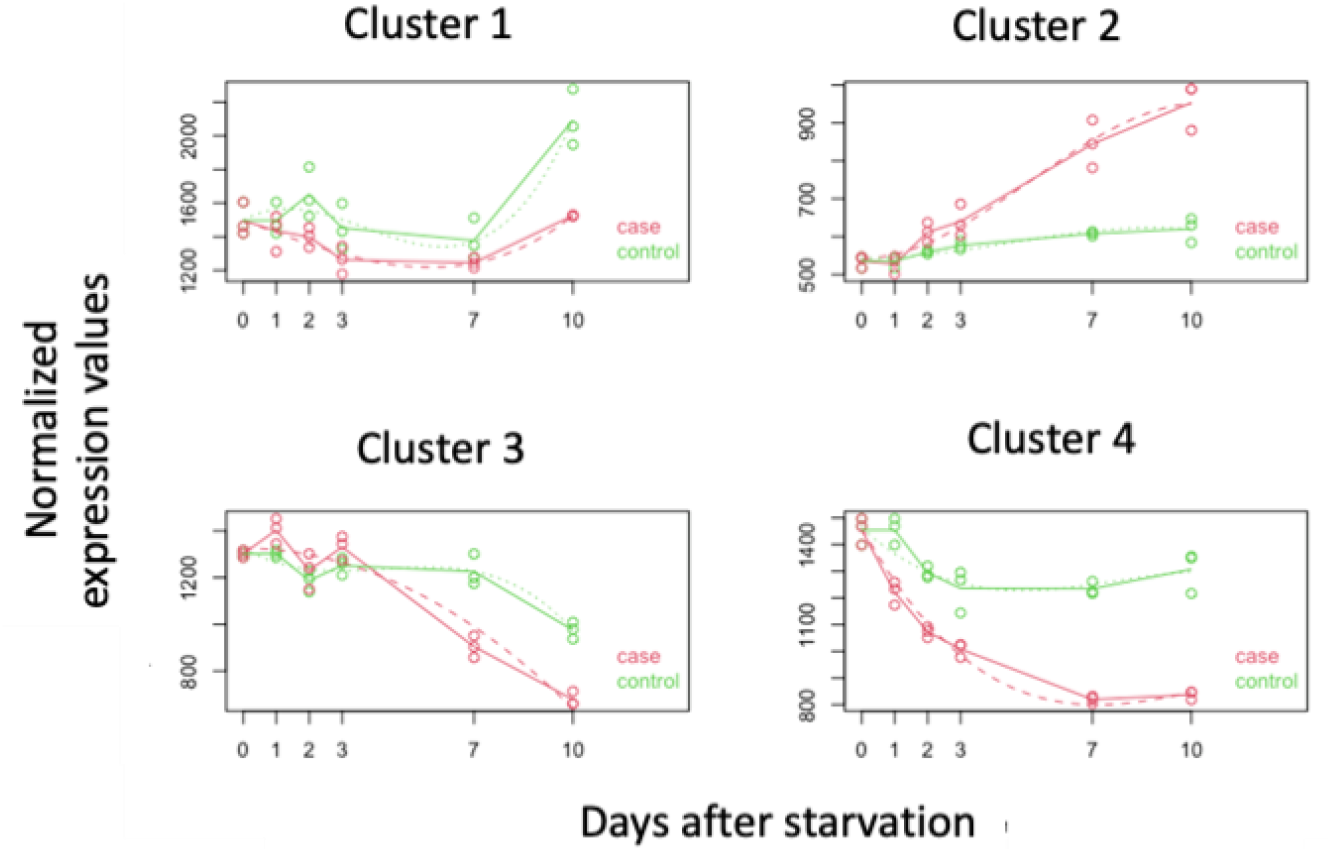
Expression profile of genes experiencing significant response to nitrogen starvation in *A. agrestis.* Each plot corresponds to a cluster of genes showing similar expression profiles over the time-course. Dots correspond to the average normalized expression values over all genes of a cluster. The three dots for each time point represent the three biological replicates. Gene expressions in the nitrogen starved (in red, case) and control samples (in green, control) are shown. Solid lines connect median expression values per time point. Dashed lines are fitted regression curves with a maximum polynomial degree of 3.

**Figure 4.**
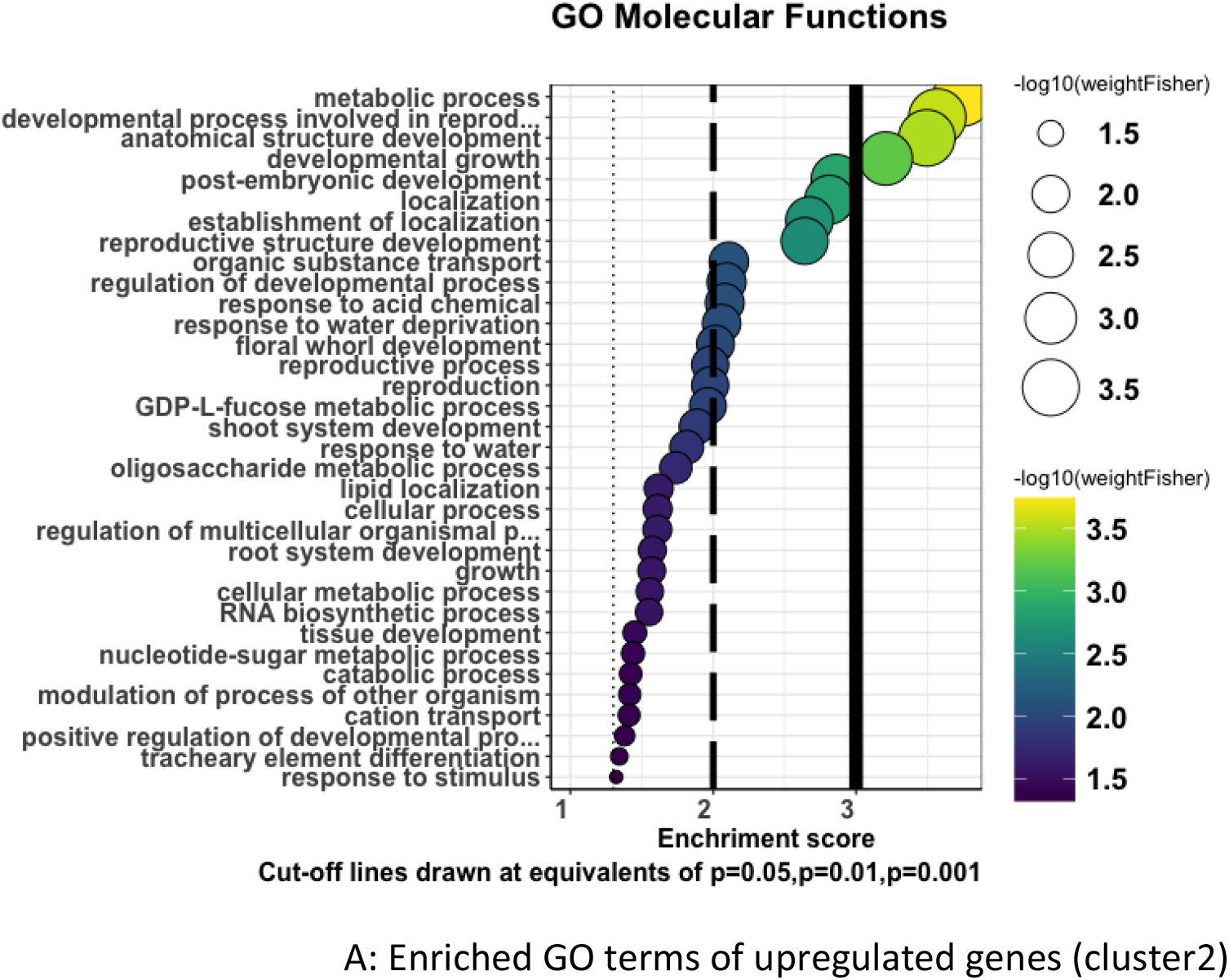
Expression profile of genes experiencing significant response to nitrogen starvation in *B. pusilla*. Each plot corresponds to a cluster of genes showing similar expression profiles over the time-course. Dots correspond to the average normalized expression values over all genes of a cluster. The three dots for each time point represent the three biological replicates. Gene expression in the nitrogen starved (in red, case) and control (in green, control) samples are shown. Solid lines connect median expression values per time point. Dashed lines are fitted regression curves with a maximum polynomial degree of 3.

### *A. agrestis* and *B. pusilla* show similar transcriptional responses to nitrogen starvation

To learn more about the dynamics of gene expression change over the time course, we identified genes showing similar temporal expression dynamics. We ran K-means clustering using a range of K values (1-9) to find a subjective optimal K value. After visual inspection of group expression profiles, we decided to use K=4 which provided a good balance between cluster distinctness and within-cluster uniformity (see results for K=2-9 in Fig. S1-S2). We did so for the two species separately.

### Anthoceros agrestis

In the hornwort *A. agrestis*, we found four major groups of genes showing well distinguishable expression dynamics (see Table S12) and the enriched GO terms for all four clusters are displayed in Table S15, Fig. S3-S6 and the enriched go terms for up/downregulated genes are shown in Fig. 5. Fig. 5A shows the enriched GO terms for upregulated genes of cluster 2 and Fig. 5B for clusters 1,3 and 4. 657 genes (cluster 2) showed gradually increasing expression under nitrogen starvation while their expression remained constant during the whole-time course in the control plants. We refer to this group as **(I) early acting upregulated genes**. GO, KEGG, and manual annotation indicated that this group was strongly enriched for genes involved in transmembrane transport both at the plasma membrane and that of the vacuole including amino-acid transporters and various ion transporters (including ammonium, nitrate/nitrite, potassium, sulfate, iron transporters). We further found that the group of **early acting upregulated genes** was enriched for lipid synthesis related genes. Besides that, we observed the upregulation of genes related to plant hormonal activity. In particular, we found that genes involved in the synthesis of gibberellic acid precursors (*KO, KAO, CPS/KS*), and strigolactones (*CCD7, CCD8*), modifying the concentration and effect of cytokinin (cytokinin receptor), signaling ABA (*NCED, abi5, bZIP*) and jasmonate action (*JAZ/Tify*) or involved with auxin concentration change were upregulated (*AUX1/LAX*). In parallel with this we also observed that genes involved in protecting the plant from oxidative damage were highly upregulated (flavonoid synthesis, phenylpropanoid pathway, polyphenol oxidases, alcohol dehydrogenases, glutathione). Nitrogen starvation also led to strong changes in carbohydrate metabolism including the upregulation of genes synthesizing trehalose and catalyzing some reactions of glycolysis and gluconeogenesis. Starvation further led to the induction of enzymes catalyzing the release of nitrogen from amino acids and enabling the assimilation of ammonium (glutamine synthase). The upregulation of various Ca-dependent kinases suggested the involvement of calcium signaling in the response to nitrogen starvation. In addition to that, we observed the expression of enzymes involved in cell wall remodeling. Finally, the early reacting gene set also contained several transcription factor (TF) genes that are likely orchestrating the molecular response (*GAPR_G2*-like [*NIN*-like], *MYB*-related, *bHLH*, *HD-III-ZIP, WRKY, RAM1* and other *GRAS, NAC, PHD, C2H2, bZIP, ARID, SET, RWP-RK, OFP, JAZ/TIFY, C2C2-Dof*).

**Figure 5.**
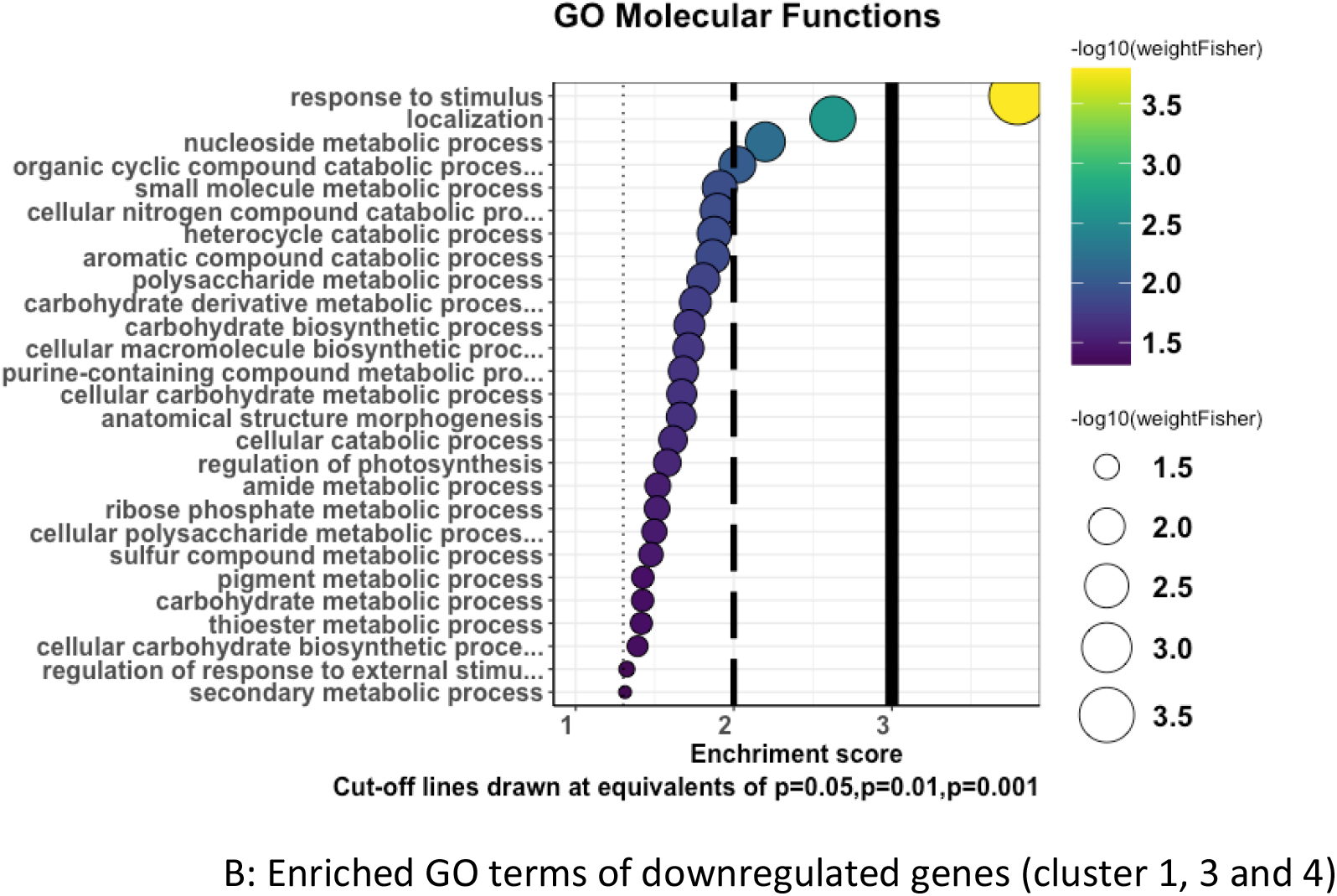
GO enrichment analysis (Molecular function ontology, P value < 0.05, Fisher’s exact test) of genes up/downregulated under nitrogen starvation in *A. agrestis*. A: enriched GO terms of upregulated genes from cluster 2. B: enriched GO terms of downregulated genes from cluster 1, 3 and 4.

A second set of genes (325 genes, cluster 4) experienced downregulation immediately after starvation, while their expression values were only slightly downregulated in the control samples. We refer to these genes as **(II) early acting downregulated genes**. These sets of genes are primarily enriched for pathways related to ribosome assembly/genesis, mRNA and rRNA processes plus amino acid synthesis. That is, genes responsible for the assembly of ribosomes and protein synthesis are immediately blocked by nitrogen starvation. We also found some TF genes being downregulated (*AP2/EREBP, GNAT, AS2/LOB, C2H2, HD-ZIP*).

Genes of cluster 3 were also downregulated but their expression decrease started only after the 3^rd^ – 4^th^ day of starvation. We refer to these genes as **(III) later acting downregulated genes**. This gene set was primarily enriched for photosynthesis-related genes, such as photosynthetic complexes (PSI, PSII) and genes responsible for chlorophyll synthesis suggesting that blocking of photosynthetic processes occurs somewhat later during nitrogen starvation. Some genes related to hormonal response and/or transcription factors were also members of this group like cytokinin oxidases (*CKX*), a *YUCCA* gene, ABA and cytokinin synthesis genes (*ABA4, IPT*), *PLATZ, C3H*.

Finally, genes of cluster 1 showed relatively rapid but weak downregulation during the first three days of starvation and their expression only slightly recovered after seven to ten days. Functionally, this cluster showed similarity to the **later acting downregulated genes** (cluster 3) because it contained genes involved with ribosome biogenesis, but also in protein synthesis, protein translation, proteasome building, and protein folding.

### Blasia pusilla

The temporal dynamic of *B. pusilla* genes reacting to nitrogen starvation was similar to those of *A. agrestis* (Fig. 6A and B, and see Table S14, GO enrichment results in Table S16 and Fig. S7-S10). Nevertheless, genes of cluster 2 showed a more gradual expression increase and reached their maximum expression value after 10 days of starvation, while genes of cluster 4 peaked already by the third day. We further refer to these two clusters as **gradually** and **rapidly reacting upregulated genes**. The **gradually reacting** group contained genes increasing nitrogen and sulfate uptake from the environment and their assimilation within the cell (glutamine synthase, ammonium, nitrate/nitrite and oligopeptide transporters). We also detected that this group contained several hormonal response genes related to stress response such as auxin (*SAUR*), *ABA* (*ABF*), gibberellin precursors (*KAO*) and cytokinin (*CYP735A*). Several genes related to cell signaling and protection against oxidative damage were upregulated such as phosphatidyl inositol and flavonoid synthesis genes, polyphenol oxidases and peroxidases. There was an indication of intensified signaling, such as increased expression of various kinases including Ca-dependent kinases suggesting that starvation reaction may be linked to Ca signaling. The increased activity of histone acetylase genes suggests that epigenetic reprogramming is taking place and may contribute to transcriptional reorganization. Finally, we also found that multiple TF genes were also part of this group (*AS2/LOB, bHLH, AP2/EREBP*).

**Figure 6.**
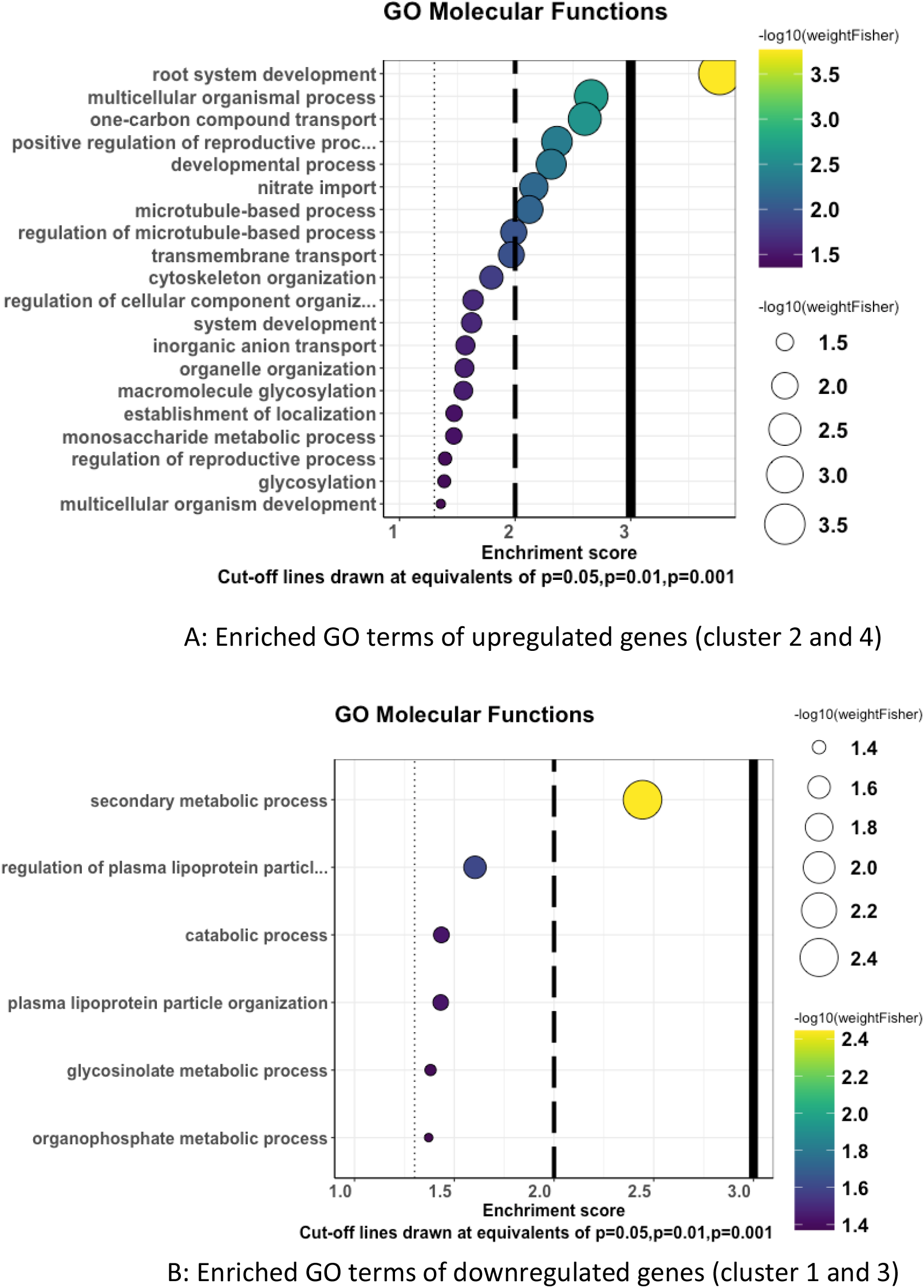
GO enrichment analysis (Molecular function ontology, P value < 0.05, Fisher’s exact test) of genes upregulated (genes of cluster 2 and 4) during nitrogen starvation in *B. pusilla*. A: enriched GO terms of upregulated genes from cluster 2 and 4. B: enriched GO terms of downregulated genes from cluster 1 and 3.

The group of **rapidly reacting upregulated** genes contained genes with similar functions. We observed multiple genes involved with nitrogen uptake, such as ammonium and nitrite/nitrate transporters. Besides the previously mentioned transporters, we also detected the immediate upregulation of sulfate transporters. An exclusive set of genes not detected in the gradually upregulated group were those involved with cell wall modifications/remodeling. Similarly, multiple mRNA, protein and amino acid synthesis-related genes were present in the rapidly reacting but not in the gradually upregulated group of genes. Finally, we also found a handful of TF genes that reacted rapidly to nitrogen starvation (*AP2/EREBP, Trihelix, bHLH, TRAF, NAC, MYB-related, ZF-HD, bZIP*).

Downregulated genes were also distributed in two clusters. Genes of cluster 1 were rapidly downregulated after nitrogen starvation, reaching their minimum expression level already on day 3. These genes are referred to as **early acting downregulated genes**. We found that multiple photosystem I/II and chlorophyll synthesis-related genes occurred in this group. Similarly, genes involved with ribosome biogenesis and translation were present as well as genes involved with signaling, auxin and gibberellin response, DNA replication and cell wall remodeling. In contrast to *A. agrestis*, we found that multiple TF genes were also part of this group (*GRAS, NAC, bHLH, GATA, AP2/EREBP, C2H2, SET, bZIP, WRKY, HD-ZIP*). Therefore, besides some TF genes, this cluster of **early downregulated genes** was functionally like cluster 2 of *A. agrestis*. While photosynthesis related genes reacted somewhat later in *A. agrestis* they did so earlier in *B. pusilla*.

Finally, genes of cluster 3 were slightly downregulated and showed relatively low expression in the nitrogen starved samples while regained expression in the control samples. Genes of this cluster are functionally overlapping with those found in the early downregulated group. For instance, we found genes involved in nitrogen preservation such as amino acid hydrolysis and amino acid synthesis. Similarly, some photosynthesis related genes were also present. Furthermore, auxin efflux carriers, signaling protein kinases and some cell wall synthesis genes were also present. Finally, we also found some TF genes to be downregulated (*NAC, C2H2, bHLH, AS2/LOB*).

### Evolutionary and functional conservation of the nitrogen starvation response between A. agrestis and B. pusilla

We hypothesized that part of the starvation response is evolutionary conserved between the liverwort and hornwort models used and could have been present in their most recent common ancestor (the common ancestor of bryophytes). We found 53 orthogroups showing shared upregulation under nitrogen starvation both in *B. pusilla* and *A. agrestis* (see Table S17, S18). GO, KEGG enrichment analysis and manual annotation of genes implies that this set of genes is enriched for plasma membrane transport processes aiding the uptake of various ions, nitrogen fixation and organic material transport. Another set of orthogroups implied that hormonal responses are also shared. We found that bZIP transcription factors (TFs) (*ABF*), cytokinin-modifying enzymes, gibberellin precursor synthesis genes, and auxin action related genes were upregulated in both species. A third set of orthogroups suggested that response of the major signal transduction and signaling cascades to nitrogen starvation is also conserved between the two species. For instance, orthogroups containing leucine-rich repeat protein kinases, protein kinases (nodule lectin) and calcium-dependent kinases show shared upregulation in both species during nitrogen starvation. We also found orthogroups containing the enzyme involved in trehalose synthesis (trehalose-6-phosphate synthase) upregulated in both species producing the essential and potent signaling molecule as well as important osmotic protectant within plant cells, trehalose. A fourth set of orthogroups suggests that nitrogen recycling from nucleotides or from chlorophyll is also evolutionary conserved between the two species. We found that orthogroups containing genes with function in obtaining nitrogen from purine nucleotides (amidase (allantoate deiminase (*allC*))) or orthogroups containing homolog of the stay-green gene that degrade chlorophyll are upregulated in both species. Finally, several shared upregulated orthogroups contained genes with functions either related to pathogen defense, or to protection from oxidative damage. This involved enzymes of the phenylpropanoid pathway, various peroxidases, and subtilisin-like proteases.

The 26 orthogroups exhibiting shared downregulation showed similar properties to what we have found in the separate analysis of the two species (see Table S17, S19). Downregulated genes were involved in transcription and translation processes which appeared to be blocked during nitrogen starvation. Similarly, chlorophyll synthesis and cell-wall expansion related genes and various metabolic processes were downregulated.

While the systemic response to nitrogen starvation was largely conserved between the model hornwort and the model liverwort, we also observed some notable differences. Our comprehensive TF annotation revealed that upregulation of various TF families was either specific to the hornwort, or to the liverwort. We found that some orthogroups containing *HD-ZIP*, *WRKY*, and *TIFY* TFs were only upregulated in the hornwort, while *AS2/LOB, AP2/EREBP, Trihelix* and *TRAF* were only in the liverwort (Fig. 7). Similarly, orthogroups containing strigolacton precursor enzymes and some lipid biosynthesis pathway genes were only upregulated in the hornwort model.

**Figure 7.**
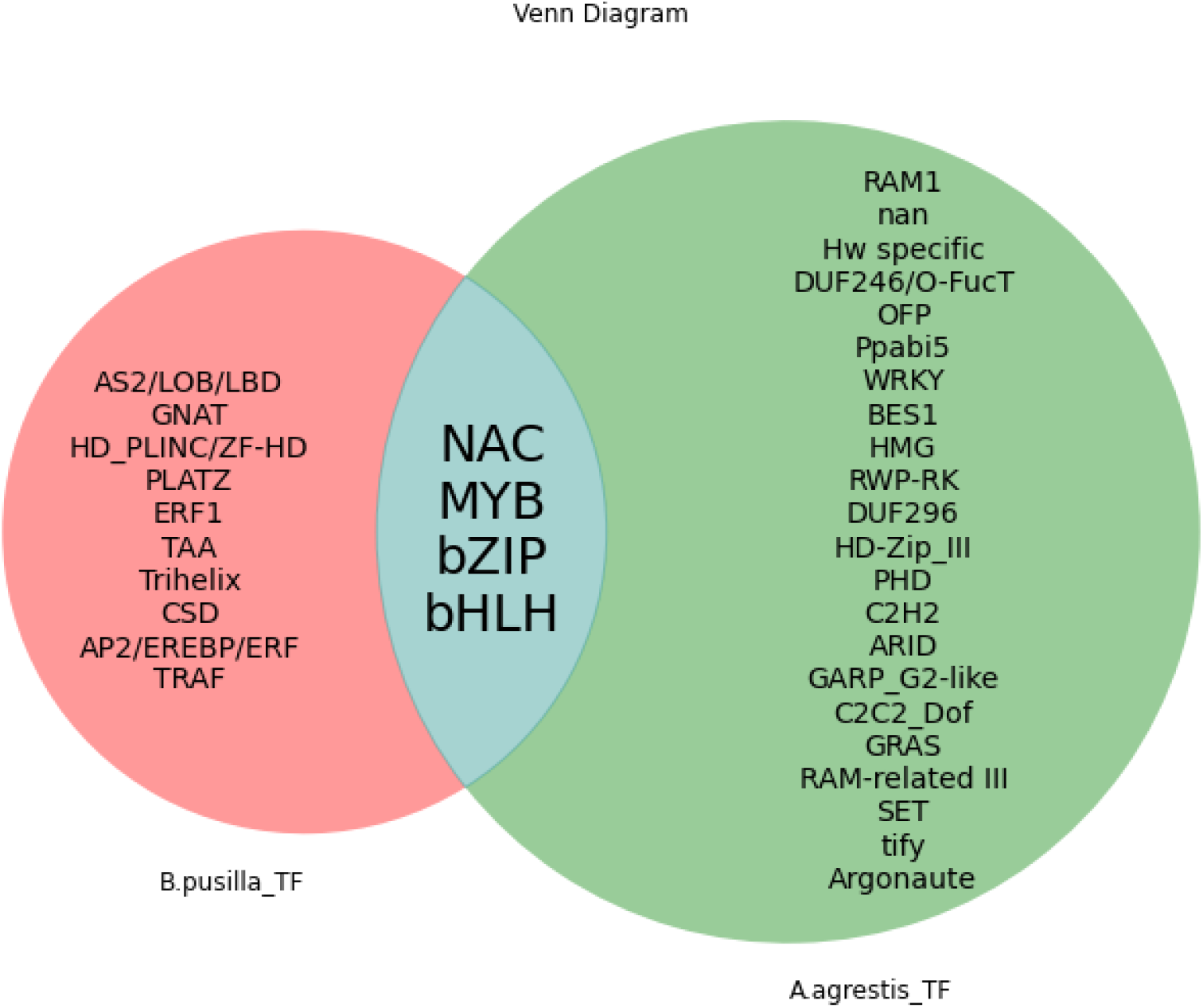
Venn diagram illustrating upregulated transcription factor families (TFs) in the liverwort (*B. pusilla)* and the hornwort (*A. agrestis)*. The intersection refers to shared upregulation.

### Conservation of the response to nitrogen starvation among bryophytes and across land plants

We hypothesized that all land plants deeply share a starvation response which was further extended with symbiont and lineage-specific modules to regulate symbiotic interactions between plants and their microbes. Both the hornwort *A. agrestis* and the liverwort *B. pusilla* can establish intimate symbiotic interaction with cyanobacteria while such interactions are not observed in other liverworts such as *M. polymorpha* and *M. paleacea*. Therefore, we reasoned that all four liverworts are expected to share a common response to nutrient starvation which has been extended with further modules in the hornwort (*A. agrestis*) and liverwort (*B. pusilla*) to enable a more intimate interaction with cyanobacteria. We identified 21 orthogroups showing shared upregulation in *A. agrestis* and *B. pusilla* during nutrient starvation but no response in *M. paleacea* and *M. polymorpha* (See Table S20-S23). These 21 orthogroups contained genes with putative functions including beta-glucanases, molybdate transporters, aldehyde dehydrogenases, trehalose synthetases, and urea transporters. We also found that several orthogroups containing TFs showed shared upregulation among others, bZIPs (*ABF*).

We further reasoned that if upregulation of these orthogroups is required to specifically attract cyanobacteria, they should not be upregulated in vascular plants establishing symbiotic interactions with either AMF or nodule-forming bacteria but not with cyanobacteria. Nevertheless, we found that most of the 21 orthogroups were also upregulated in one or more vascular plants not able to establish intimate interaction with cyanobacteria (Fig. 8).

**Figure 8.**
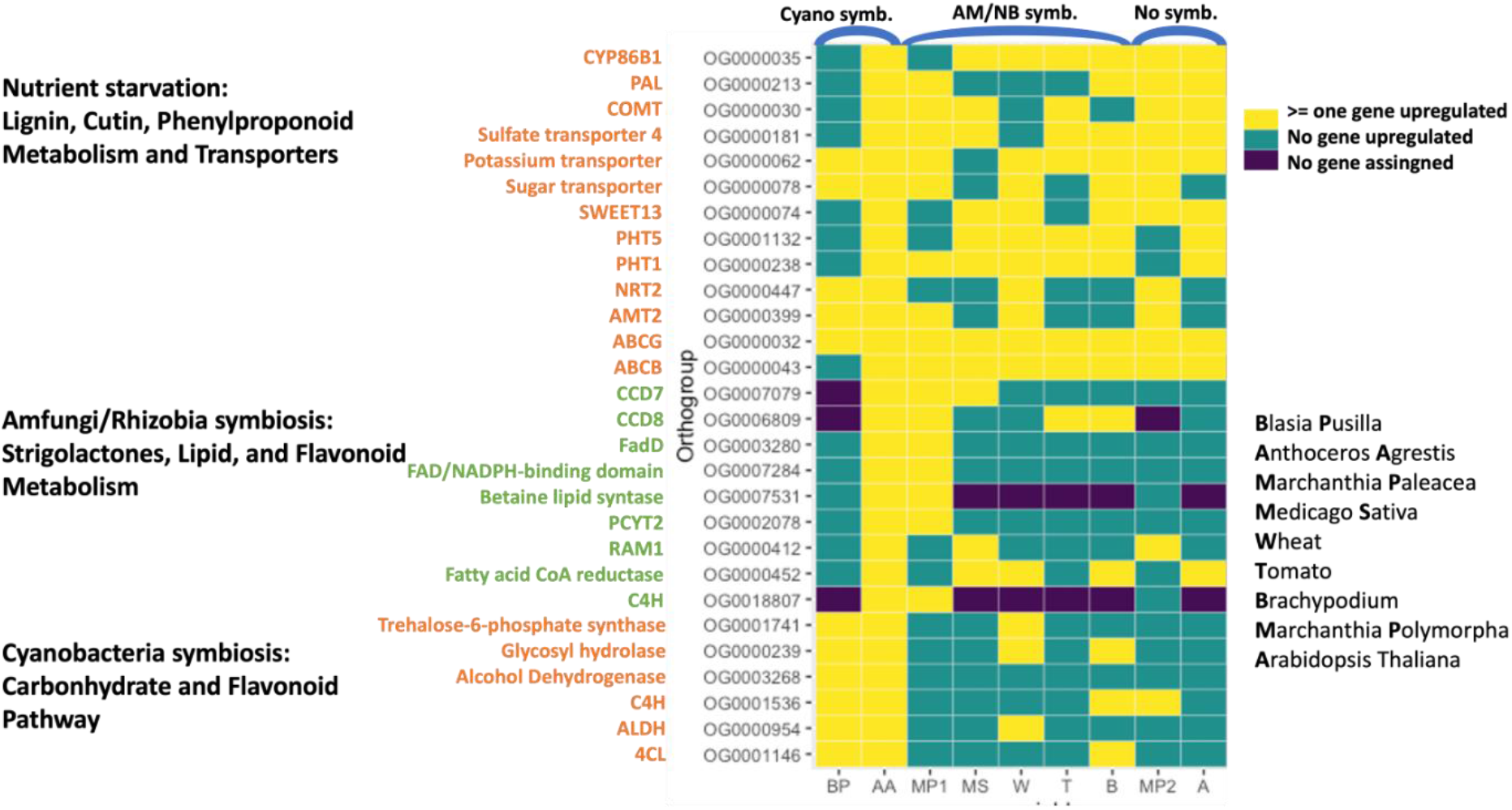
Selected orthogroups upregulated under nutrient starvation (either nitrogen or phosphate starvation) a wide range of symbiotic and non-symbiotic vascular and non-vascular plant species (cyanobacteria-symbiosis, AM fungi symbiosis, nodule-forming bacteria symbiosis, and non-symbiotic species). Black: no gene is assigned to this orthogroup, green: orthogroup contains genes but none of them are upregulated, yellow: at least 1 gene assigned to the orthogroup is upregulated in response to starvation.

## Discussion

It has been proposed that the regulatory network responsible for initiating plant-microbe symbiotic interactions has evolved from a conserved, ancient nutrient starvation response that was already present in the common ancestor of all land plants (Isidra-Arellano et al., 2021; Paries and Gutjahr, 2023). Later this network has further recruited and extended to genes specifically regulating the intimate interaction between plants and AM fungi, nodule-forming bacteria and potentially cyanobacteria (Ma and Chen, 2021; Delaux and Schornack, 2021; Puginier et al., 2022; Shi et al., 2022). Here we investigated this hypothesis by exploring gene expression changes during nitrogen starvation in the two major lineages of bryophytes, hornworts and liverworts, able to establish intimate symbiotic interaction with cyanobacteria (Meeks, 1998; Álvarez et al., 2023a). By conducting comparative analyses with symbiotic and non-symbiotic vascular and non-vascular plants, we found that the systemic gene expression response to nitrogen/phosphate starvation is similar in bryophytes and vascular plants. Additionally, examination of the dataset allowed us to identify several groups of compounds that may serve as hormogonia-inducing factors (HIF), while providing little evidence for an extensive set of genes involved in priming the host plant for the cyanobiont. These findings allow us to propose a hypothetical description of the starvation response and its convergence and divergence in the hornwort and liverwort models, which is detailed below (Fig. 9).

**Figure 9.**
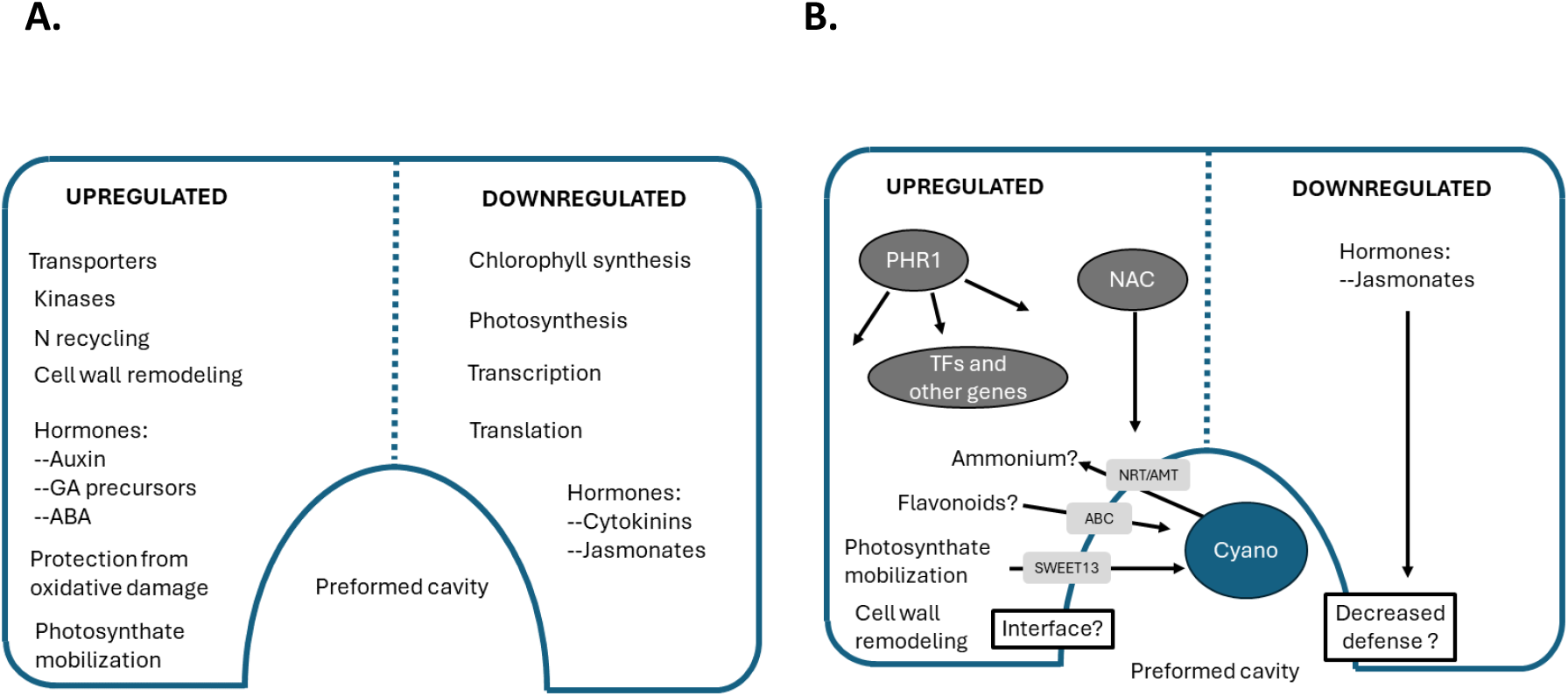
Systemic response to nitrogen starvation and potential priming of the host to the cyanobiont. A.) Up- and downregulated biological processes under nitrogen starvation shared by *B. pusilla* and *A. agrestis*. B.) Biological processes/genes potentially contribute to priming the plant host to the cyanobiont. Transporters are shown on light gray background, transcription factors on dark gray background in white letters.

### Systemic response to nitrogen starvation appears to be conserved across bryophytes and vascular plants

Response to nitrogen starvation in vascular plants involves sensing nitrogen levels, transduction of the signal into the cell, and adjusting gene expression and metabolism to preserve metabolic balance (Varala et al., 2018; Gaudinier et al., 2018; Wang et al., 2019; Ueda and Yanagisawa, 2019; Ueda et al., 2020). We observed conservation in the response of *A. agrestis* and *B. pusilla* as well as with vascular plants in most of these steps.

#### Sensing and signaling of low nitrogen environment

Low nitrogen levels are sensed by *NRT* genes (nitrate sensor) which work together with calcium channels and activate calcium signaling pathways inducing transcription factors as well as further genes involved in nitrogen uptake, nitrogen metabolism and auxin regulation (Mascia et al., 2019; Dong et al., 2022; Fang et al., 2022; Adavi and Sathee, 2023). We observed that major components of the calcium signaling pathway (calmodulins, calcium-dependent protein kinases, serine-threonine protein kinases) were induced both in *A. agrestis* and *B. pusilla* during starvation. This suggests that calcium signaling is a common theme between nitrogen starvation in bryophytes and vascular plants. Calcium signaling is also central to transmit systematic response to various stresses (Dong et al., 2022). Whether calcium signaling involves calcium spiking or other mechanisms is unknown.

#### Nitrogen Uptake, Transport, assimilation, and Related Modulators

In vascular plants nitrogen limitation rapidly activates nitrogen uptake systems enabling its transport from the environment to the cell (Fan et al., 2017; Aluko et al., 2023). Nitrate (*NRT*), ammonium (*AMT*) transporters are upregulated to increase nitrogen uptake and assimilation (Rui et al., 2022; Aluko et al., 2023). In vascular plants, activation of nitrogen uptake and assimilation mainly occurs in the root. While roots are not present in non-vascular plants, rhizoid plays an analogous role (Jones and Dolan, 2012; Szövényi et al., 2019). We found that nitrate and ammonium transporter homologs were upregulated in the *B. pusilla* and *A. agretis* systems like in vascular plants. Specifically, an ammonium transporter *AMT2* and a nitrate transporter *NRT2.2* homolog were upregulated both in *A. agrestis* and *B. pusilla*. Amino acid and peptide transporters were also upregulated that likely help the uptake of amino acids/peptides from the environment. In vascular plants nitrogen is mainly available and taken up as NO_3_^-^, and NH_4_^+^ (Masclaux Daubresse et al., 2010). NO_3_^-^ is then reduced to NO_2_^-^ by the nitrate reductase (*NR*) and then transported to the chloroplast together with NH_4_^+^. NO_2_^-^ is reduced to NH_4_^+^ by the nitrite reductase and together with the transported NH_4_^+^ fed into the GS/GOGAT cycle to be assimilated into glutamate which serves as a precursor of multiple metabolites (Dubey et al., 2014). When vascular plants are starved of nitrogen, protein reserves of the chloroplast are proteolyzed and NH_4_^+^ and amino acids are recycled into glutamine by GS. We found that nitrogen starvation led to the upregulation of the *GS* and not the *GOGAT* enzyme in both species *B. pusilla* and *A. agrestis* suggesting recycling of protein resources to maintain nitrogen balance of the cell. This implies the presence of a physiological response conserved between vascular plants and bryophytes in preserving nitrogen balance.

Besides nitrogen transporters, our transcriptome analyses also suggested that ABC transporters are induced under nitrogen starvation (both *ABCG* and *ABCB* transporters). *ABCG* transporters transport lipids and or (iso)flavonoids as well as phytohormones. Even though it has been shown that cyanobacteria receive carbohydrates (at least some of it in the form of sucrose) from its plant partner (Kaplan and Peters,1998), the necessary transporters of either symbiont have not been identified. We found that *SWEET13* homologs were upregulated in *A. agrestis* under nitrogen starvation conditions likely transporting sugars towards the cyanobiont.

#### Hormonal response

In flowering plants, nitrogen and phosphate starvation cause hormone synthesis activation/repression. Hormonal changes lead to both physiological and morphological adaptations increasing nutrient usage. Auxin synthesis and transport are well known to be activated under low nitrogen conditions to change root shoot allocation (Sultana et al., 2020). The most obvious effect of auxin is to trigger root hair growth under nitrogen and phosphate limitation (López-Bucio et al., 2002). Auxin response is apparent in both species data sets (*GH3, AUX1*) and likely trigger rhizoid development as was shown in the liverwort *M. polymorpha* (Rico-Reséndiz et al., 2020; Rico-Resendiz et al., 2021; Suzuki et al., 2021). Because auxin is necessary for normal organogenesis in *M. polymorpha*, it is possible that it also plays a role in the developmental process of the cavities containing the cyanobacterial cells in both *B. pusilla* and *A. agrestis* (Suzuki et al., 2023).

Similarly, a low nitrogen environment is known to promote ABA accumulation which will affect lateral root formation, nitrate sensing and transport in vascular plants (Ondzighi-Assoume et al., 2016; Wang et al., 2018; Lee et al., 2020). Overall, ABA helps to increase tolerance to low nitrogen conditions in general. We assume that ABA has a similar stress alleviating effect in the hornwort and the liverwort, however its effect awaits further scrutiny.

The role of cytokinin in nitrogen starvation of vascular plants is poorly explored but phosphate starvation reduces cytokinin levels in *A. thaliana* thereby affecting the shoot to root ratio and enabling root hair growth (Kiba et al., 2011; Lv et al., 2021). Similarly, it was found that cytokinin oxidase genes, decreasing the amount of active cytokinin, are upregulated during phosphate starvation in *M. polymorpha* (Rico-Resendiz et al., 2020). Furthermore, cytokinin is known to promote rhizoid development in *M. polymorpha* like its effect on root hair growth in flowering plants and may have a similar function in *A. agrestis* and *B. pusilla* (Rico-Reséndiz et al., 2020; Rico-Resendiz et al., 2021).

GA levels increase in vascular plants under nitrogen starvation, and it likely triggers stress tolerance against low nitrogen conditions. GA receptors and canonical gibberellins are not present in bryophytes (Guillory and Bonhomme, 2021), but hormonal diterpenoids regulate various aspects of development in mosses as well as in liverworts. For instance, *DELLA* proteins promote protection against oxidative stress response and coordinate growth across various environmental inputs in the liverwort *M. polymorpha* (Hernández-García et al., 2021). Our results suggest that this response is also likely conserved between liverworts, hornworts, mosses and vascular plants.

Jasmonates are also upregulated under defense in flowering plants and are likely to regulate sugar, amino acid synthesis and NH_4_^+^ uptake (Sun et al., 2020; Lv et al., 2021). By contrast, in the hornwort model some *JAZ* homologs were upregulated which are known to repress jasmonate synthesis in liverworts and promote cell growth and reproductive fitness (Monte et al., 2019). Furthermore, reduced jasmonate synthesis leads to repressed defense responses. While the role of *JAZ* upregulation during nitrogen starvation in the hornwort is currently unclear, it is possible that it decreases plant defenses against pathogens and this way triggers the priming of the host plant to the cyanobiont (Monte et al., 2019).

Together with jasmonates, salicylic acid (SA) also plays an important role in the defense against pathogens in vascular plants (Ding and Ding, 2020). SA has been proposed to be involved in the signaling between host plant and the cyanobiont in the *Azolla-Nostoc* symbiosis (de Vries et al., 2018, 2023). While most biosynthetic enzymes of SA are present in the hornwort genome, we found none of them upregulated during nitrogen starvation. Therefore, our data do not support the findings observed in the *Azolla-Nostoc* symbiosis.

#### Transcription factors

The nitrogen starvation induced calcium signaling pathway converges on various transcription factor genes that act as master regulators of the nitrogen starvation response. In vascular plants *bZIP* transcription factors are usually upregulated and transmit ABA response, increase nitrogen uptake and regulate root branching (Zhang et al., 2015; Yang et al., 2019; Ma et al., 2020). *WRKY* transcription factors are also induced and increase nitrogen stress tolerance (Curci et al., 2017; Wang et al., 2019). Upregulation of *MYB* and MYB-related transcription factors increase nitrogen influx, development, cell cycle and morphology (*NIGT*) (Curci et al., 2017; Wang et al., 2019), while CCAAT-binding TFs (such as *TaNFYA)* are important regulators of nitrate transporters (Qu et al., 2015; He et al., 2015). *C2H2* TFs are known to trigger low-nitrogen adaptation and *Dof* TFs can induce ammonium transporters as well as the synthesis of carbonic scaffolds to be used for nitrogen assimilation (Yanagisawa et al., 2004; Noguero et al., 2013; Chen et al., 2017).

We found that multiple members of these TF families were also upregulated in either one or both species investigated. For instance, *bZIP* TFs of the ABA responsive element binding factor (*ABF*), *MYB-related*, *bHLH* and *NAC* TFs were upregulated during nitrogen starvation in both species. Homologs of these TFs were also upregulated under nitrogen starvation in *M. polymorpha* and phosphate starvation in *M. paleacea* (Radhakrishnan et al., 2020; Rico-Reséndiz et al., 2020). Therefore, these TFs appear to represent a conserved subset of master regulators that could have been already present in the common ancestor of most bryophytes and some of them may have master regulatory functions even shared with vascular plants.

*NAC* TFs are also often upregulated under nitrogen starvation and affect root growth, branching and nitrate uptake (Vidal *et al*., 2013, 2014; He *et al*., 2015). Stress induced *NAC* TFs have recently also been found to be induced under nitrogen starvation in rice (Qi et al., 2023). Furthermore, it was also shown that the *OsNAC1* gene directly induced the expression of multiple nitrate transporters (*NRT* genes) and led to higher nitrogen use (NUE) efficiency, biomass and grain yield (Tang et al., 2019; Li et al., 2020a; Qi et al., 2023). Therefore, we hypothesize that the connection between nitrogen starvation induced *NAC* TFs and *NRT* genes is likely to be conserved among liverworts, hornworts and vascular plants. Future work should reveal to what extent this conserved systemic regulatory module is linked to further modules potentially enabling colonization of the host plant by the cyanobiont.

MYB-related TFs, especially phosphate response 1 (*PHR1*), is a master regulator of the phosphate starvation response in flowering plants as well as in *M. polymorpha* (Rouached et al., 2010; Rico-Reséndiz et al., 2020; Isidra-Arellano et al., 2021; Tan et al., 2023; Paries and Gutjahr, 2023). Furthermore, *PHR1* is essential in priming the host plant for plant AMF and rhizobia symbiosis by regulating plant immunity and a diverse set of genes necessary for the initiation and establishment of the symbiotic interaction (Isidra-Arellano et al., 2021; Paries and Gutjahr, 2023). Importantly, our analysis indicates that the upregulated Myb-related transcription factors are homologous to the *PHR1* gene of *A. thaliana* and *M. polymorpha*. Therefore, it is likely that the upregulation of *PHR1* homologs and their master regulatory function in nitrogen/phosphate starvation are shared between the hornwort, the liverwort models (and other liverworts) and vascular plants. How much the regulated gene set differs between flowering plants and our two model species needs further investigation.

Multiple *bHLH* TFs are known to be upregulated under various stresses in flowering plants including phosphate and nitrogen-starvation (Yang et al., 2016; Subudhi et al., 2020). Some of these genes regulate root growth while some others influence phosphate/nitrogen starvation response genes (Crombez et al., 2019; He et al., 2021). Phylogenetic analysis shows that the *bHLH* gene identified in the hornwort and liverwort models forms a monophyletic group with *Azolla filiculoides* genes. Furthermore, it is embedded in a large clade containing only bryophyte *bHLH*s. Therefore, this gene could potentially be important for the cyano-hornwort/liverwort symbiotic interaction.

While some TFs reacted similarly in the hornwort and liverwort species (*A. agrestis* and *B. pusilla*) investigated, many TF family genes showed divergent expression patterns whose potential function remains to be investigated. For instance, *AS2/LOB* TFs are known to repress nitrate starvation response genes in *A. thaliana*, while they were upregulated only in our model liverwort *B. pusilla* (Rubin et al., 2009; Dash et al., 2015). Similarly, *AP2/EREPB* TFs are known to be induced in nitrogen starved flowering plants (Yang et al., 2015), but they were only upregulated in the liverwort model studied. Finally, the role of *Trihelix* and *TRAF* TFs in nitrogen starvation response of *B. pusilla* is unclear.

Intriguingly, we found many more TF genes to be upregulated under nitrogen starvation in the hornwort than in the liverwort. Some of them are well-known to show similar responses in flowering plants including, *HD-ZIP III* (Lin et al., 2022), *WRKY* (Wang et al., 2019), *MYB* (Zhang et al., 2012; Wang et al., 2019, 2020b), *C2H2* (SUN et al., 2012; Chen et al., 2017), *C2C2-Dof* (Yanagisawa et al., 2004; Curci et al., 2017), *GARP* (Medici et al., 2015) and *RWP_RK* (Chardin et al., 2014; Wu et al., 2020) TFs. It is unclear why these TFs are not responsive in the liverwort *B. pusilla* although these genes are mostly present in the genome of liverwort *B. pusilla.* We speculate that the difference between the hornwort and the liverwort model may be partly due to their divergent habit/growth form and thallus organization as well as their ability to engage in symbiotic interaction with AMF (see below).

### Nitrogen starvation and symbiotic interaction

It has been known for a long time that nutrient starvation (Pi/N) is a prerequisite for the establishment of plant-AMF and plant-rhizobia symbioses (Ma and Chen, 2021; Das et al., 2022). In these interactions master regulators of the phosphate (*PHR*) or nitrogen starvation syndrome (*NIN*) directly regulate genes involved in pre-colonization signaling of the symbiotic interaction with AMF or rhizobia, respectively (Diédhiou and Diouf, 2018; Luo et al., 2023). As the very first step of symbiosis initiation, these master regulators activate the synthesis and secretion of special chemical signals, strigolactones in AMF and flavonoids in nodule-forming symbiosis, which trigger the growth and colonization potential of the microbes. As a response to the chemical signals, the microbes release signaling molecules perceived by host plant receptors and activate precolonization signaling (Oldroyd, 2013; Zipfel and Oldroyd, 2017; Oldroyd and Leyser, 2020; Jhu and Oldroyd, 2023).

Similarly, plant-cyanobacteria symbiotic interactions are also suppressed in the presence of sufficient amounts of combined nitrogen (Meeks, 1998; Adams and Duggan, 2008; Álvarez et al., 2023b). Starved host plants release the so-called hormogonia inducing factor (HIF) triggering the conversion of vegetative cyanobacterial filaments to motile hormogonia that can colonize the host plant (Campbell and Meeks, 1989; Bergman et al., 1996; Meeks, 2003; Adams, 2005). Nevertheless, the exact chemical composition of the HIF is poorly understood and appears to vary across species. Previous studies found that monosaccharides (arabinose, glucose), diacylglycerols, or even various pigments likely function as HIFs (Rasmussen, 1994; Nilsson et al., 2006; Adams and Duggan, 2012; Liaimer et al., 2015; Hashidoko et al., 2019; Alvarenga et al., 2022; Carrell et al., 2022). Intriguingly, our transcriptomic analysis suggests that genes responsible for the synthesis of some of these compounds are upregulated under nitrogen starvation both in the hornwort and in the liverwort system investigated. Furthermore, they are rarely upregulated in species forming no symbiotic interaction with cyanobacteria.

For instance, we found that genes involved in flavonoid synthesis as well as *ABCG* transporter homologs known to transport isoflavonoids are upregulated in both the liverwort and the hornwort systems. This suggests that flavonoid type molecules can potentially function as HIF components. This is in line with previous findings that flavonoids can affect the activity of a gene cluster involved in hormogonia formation of the cyanobiont (Cohen and Yamasaki, 2000). Hormogonia are the motile forms of cyanobacteria necessary for the colonization of the host plant. Similarly, we observed the upregulation of trehalose-synthetase genes. Trehalose was recently proposed to potentially be transported from the host plant to the cyanobiont and may have HIF-like activity in peat mosses (Carrell et al., 2022). Therefore, both compounds should be investigated for their HIF activity in the future.

In the plant-AMF and nodule-forming bacteria symbioses starvation is not only necessary to attract and mobilize the microbial partner but it is also essential to prime the host for the interaction by adjusting its physiology as well as creating specialized cellular structures guiding microbial colonization (Isidra-Arellano et al., 2021; Shi et al., 2021; Das et al., 2022). While specialized cellular structures (cavities) hosting the microbes are performed and do not need to be induced in the hornwort- and liverwort-cyanobacteria symbioses, it is unclear how much further priming is necessary to accept the cyanobiont. Collectively, our data suggests that some genes induced by nitrogen starvation may be involved in priming, but an extended network of genes is absent. Therefore, it is possible that no extensive priming is necessary besides chemotactic attraction and HIF-mediated mobilization of the cyanobiont. Alternatively, the absence of an extensive shared priming network may be a consequence of the independent evolution of symbiotic interaction regulation in hornworts and liverworts.

While the gene network involved in priming is unlikely to be extensive, our data provides some candidate genes whose expression is triggered by nitrogen starvation and could potentially be important in priming the plant host for symbiotic interaction. This is in line with the observations that starvation is not only necessary to attract and mobilize the cyanobiont but it is also essential to maintain symbiotic interaction (CAMPBELL and MEEKS, 1992). For instance, it is known that blocking the upregulation of nitrate transporter genes in the plant-nodule-forming bacteria interaction will inhibit the establishment of the symbiotic interaction (Wang et al., 2020a). Therefore, it is possible that the upregulation and proper localization of various transporters in the hornwort and liverwort during nitrogen starvation helps priming the host plant for the symbiotic interaction. In the hornwort and the liverwort, we also found sugar transporters (including *SWEET*) to be upregulated. This is in line with the observation that photosynthates are provided by the host plant to the cyanobiont (de Vries and de Vries, 2022). *Nostoc* with dysfunctional sugar transporters will not establish symbiotic interaction (Ekman et al., 2013), and that host plant cavities containing the cyanobiont are filled up with polysaccharide containing mucilage. Therefore, it is possible that upregulation of sugar transporters is necessary for the initiation and establishment of the symbiotic interaction by attracting, feeding and retaining the cyanobiont. Finally, we also detected alcohol dehydrogenases to be mainly upregulated in the hornwort and liverwort models during nitrogen starvation but not in species without intimate interaction with cyanobacteria. Alcohol dehydrogenase overexpression is often observed in pathogen colonization and was shown to support biotrophy (Pathuri et al., 2011). Therefore, overexpression of such enzymes may be important in priming the host plant for symbiotic interaction.

### Several nitrogen-starvation induced genes in A. agrestis are likely involved in priming the host plant for AMF symbiosis

While the above-mentioned transcriptional responses were mainly shared between the hornwort and the liverwort, a large number were specific to only one of the model species investigated. Intriguingly, more genes and TFs were upregulated in the hornwort model and many of these were shared with species exhibiting symbiotic interaction with AMF. We speculate that the hornwort-specific upregulation of these genes reflects the ability of *A. agrestis* to form symbiotic interactions with AMF.

NIN-like TFs (*NLP* genes, *RPW-RK* transcription factors) are especially interesting in this respect because they are master regulators of the nitrogen-starvation response regulating many other genes (Chardin et al., 2014). We found that multiple *RPW-RK* transcription factors homologous to the NIN-like proteins of *M. polymorpha* (Mu and Luo, 2019; Hsin et al., 2021) were significantly upregulated but only in *A. agrestis*. Therefore, our data suggests that the sensing and signal transduction of low nitrogen environments in *A. agrestis* likely involve Ca^2+^ signaling and NIN-like proteins in a similar way to what is described for vascular plants (Mu and Luo, 2019).

The upregulation of a *RAM1* homolog in the hornwort model during nitrogen starvation is also intriguing. Expression of *RAM1* together with other *GRAS* TFs is known to be induced by phosphate starvation in vascular plants establishing AMF symbiosis and is essential in inducing various genes required to form the infection thread and the interface between the host plant and the fungal partner (Gobbato et al., 2012; Xue et al., 2015). *RAM1* is also essential for lipid provisioning (Luginbuehl et al., 2017). *RAM1* is a member of the common symbiosis signaling pathway (CSP) genes with a complete set of homologs present in the hornwort genome (Li et al., 2020b). By contrast, most homologs of the CSP genes are absent from the *B. pusilla* genome. Furthermore, we found that no *GRAS* TFs were upregulated under nitrogen starvation in *B. pusilla*. This suggests that *GRAS* TFs may not be necessary to prime the host plant for the cyanobiont in the hornwort and liverwort model systems but could be important in priming the hornwort for AMF colonization. This is further supported by the upregulation of lipid synthesis genes in the hornwort *A. agrestis* but not in the liverwort *B. pusilla* (Rich et al., 2021).

Another prominent example is the upregulation of genes involved in the biosynthesis of strigolactone precursors during nitrogen starvation which was also recently observed by other studies (Radhakrishnan et al., 2020; Rich et al., 2021). We found that nitrogen starvation positively affects the expression of *D27* and the gene encoding the enzyme catalyzing subsequent steps of strigolactone biosynthesis – *CCD7.* A new form of strigolactone has recently been shown to affect AMF colonization in the liverwort *M. paleacea* (Kodama et al., 2022). It was also observed that some of the key strigolactone synthesis genes are missing from the genome of *M. polymorpha*, a sister species of *M. paleacea*, which is not able to establish symbiotic interaction with AMF. Therefore, the secretion of strigolactones into the environment appears to be necessary for AMF interaction. When comparing the hornwort and the liverwort proteomes, we made similar observations and found that genes crucial for the biosynthesis of strigolactones were missing from the *B. pusilla* but were present in the *A. agrestis* genome. Therefore, we hypothesize that strigolactones are unlikely to be necessary to establish the plant-cyanobiont interaction. While our data and previous observations support our hypothesis, the potential role of strigolactones and *GRAS* TFs in the hornwort-cyanobacteria interaction cannot be fully ruled out and warrants further scrutiny.

### Overall conclusions

Our study also has some important applied implications for engineering plant-cyanobacteria symbiotic interactions. The observation that almost no orthogroups were specifically upregulated in the cyanobacteria host plants suggests that little host plant priming is needed to accept the cyanobiont. Therefore, the attraction and mobilization of the cyanobiont by the HIF may be the most critical step in establishing symbiotic interaction. Our data suggest that beyond that, very little if any priming is necessary in the host plant to accept the cyanobiont. Therefore, identifying compounds with HIF activity and applying them under natural conditions may be key to attracting cyanobacteria by non-host plants.

## Supporting information

Draft_genome

Transcriptomic_reference

## Supplementary data (available at Figshare https://figshare.com/s/0acebe58f3dee5065469)

Table S1. Design of the starvation experiment (*A. agrestis* and *B. pusilla*) and the number of replicates used for RNA sequencing.

Table S2. Proteomes used in the orthofinder analysis.

Table S3. RNA-seq data used for comparative transcriptomic analyses.

Table S4. Genes showing significant (padj<0.05) response (up and down-regulation) over the course of the nitrogen starvation experiment for *M. polymorpha*.

Table S5. GO annotation of *A. agrestis* genes including the molecular function, biological process and cellular localization ontologies.

Table S6. Kegg annotation of *A. agrestis* genes.

Table S7. GO annotation of *B. pusilla* genes.

Table S8. KEGG annotation of *B. pusilla* genes.

Table S9. Predicted transcription factors and genes involved in hormone synthesis and signaling (*B. pusilla* and *A. agrestis*).

Table S10. Blobtools contamination filtering of the *B. pusilla* draft genome assembly.

Table S11 Normalized expression values for *A. agrestis* over the course of the time-series experiment.

Table S12. Results of the maSigpro analysis for *A. agrestis*.

Table S13. Normalized expression values for *B. pusilla* over the course of the time-series experiment.

Table S14. Results of the maSigpro analysis for *B.pusilla*.

Table S15. GO-enrichement analyses results for *A. agrestis* (all four clusters). Table S16. GO enrichment analysis results for *B. pusilla* (in all four clusters).

Table S17. Up- and down-regulated genes and their orthogroup assignment in *A. agrestis* and *B. pusilla*.

Table S18. Upregulated genes in the *B. pusilla* and *A. agrestis*. Table S19. Genes downregulaten in the *B. pusilla* and *A. agrestis*.

Table S20. Upregulated genes in the vascular and nonvascular species investigated.

Table S21. Genes upregulated in the selected nonvascular and vascular plants under nitrogen/phosphate starvation.

Table S22. Downregulated genes in the vascular and nonvascular species investigated.

Table S23. Genes downregulated in the selected nonvascular and vascular plants under nitrogen/phosphate starvation

## Author Contribution

PS and YY conceptualized the study. YY, GS and AN generated the primary sequence data. YY, PS and GS carried out the draft genome assembly. YY and PS carried out *de novo*-transcriptomic assembly, conducted detailed transcriptomic and comparative transcriptomic analyses. YY and PS wrote the manuscript. All co-authors revised and approved the final manuscript.

## Data Availability

Raw DNA and RNA sequencing data used in this publication was submitted to NCBI Short Read Archive (SRA) under the BioProject ID PRJNA1099914 (SRA submission SUB14374705, SUB14439959, SUB14439962). The transcriptome reference and the draft genome assemblies are available on Figshare (will be replaced by DOI once published). Supplementary tables are also available on Figshare (will be replaced by DOI once published).

## Acknowledgments

We are grateful to Elke Dittmann for providing *Blasia pusilla* and cyanobacteria cultures. We are thankful to Jessy Zhang for assistance in sequencing at the Novogene UK. We are also thankful for the S3IT team and the ScienceCloud infrastructure at the University of Zurich for providing computational resources. JH and G thank CSC IT center for science (Espoo, Finland) for computational resources and advise in Finland. We also appreciate early discussions about the topic with John Bowman and Charles A. Delwiche.

