## Supplementary material for "Nitrogen starvation response in hornworts and liverworts provides little evidence for complex priming to the cyanobiont": Supp_Figs_S1-S10

Figure S1. **Expression clusters** (**maSigpro analysis) for *A. agrestis.***

**kcluster=2.**

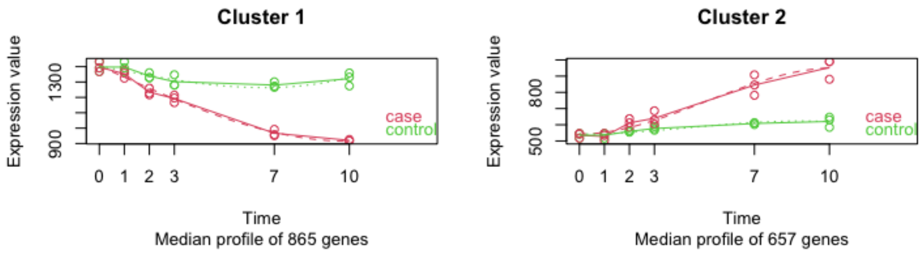


**kcluster=3.**

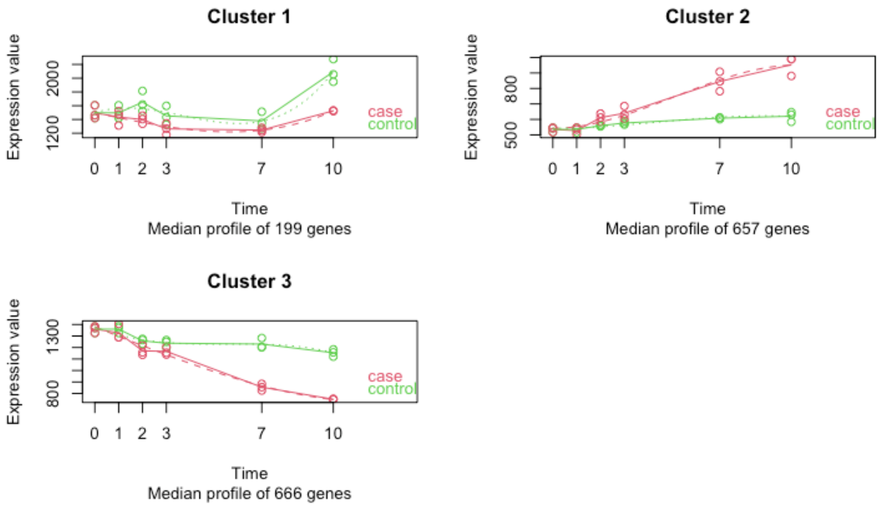


**kcluster=4.**

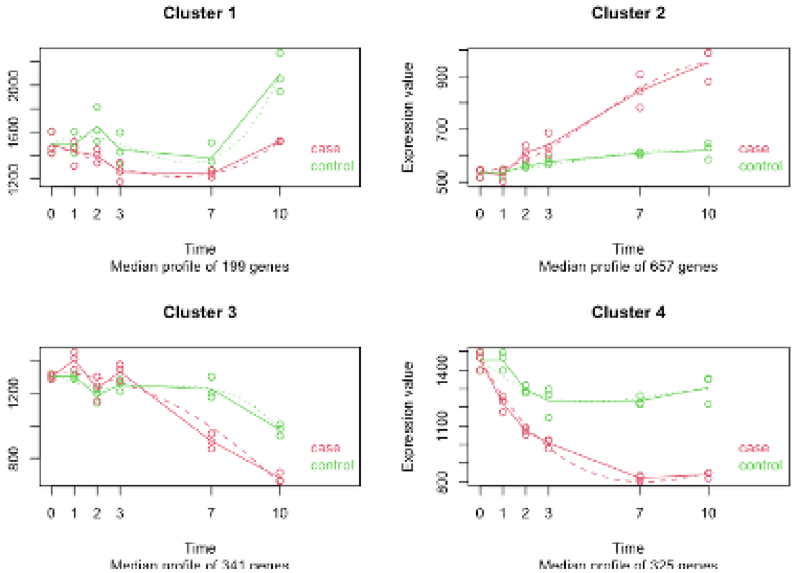


**kcluster=5.**

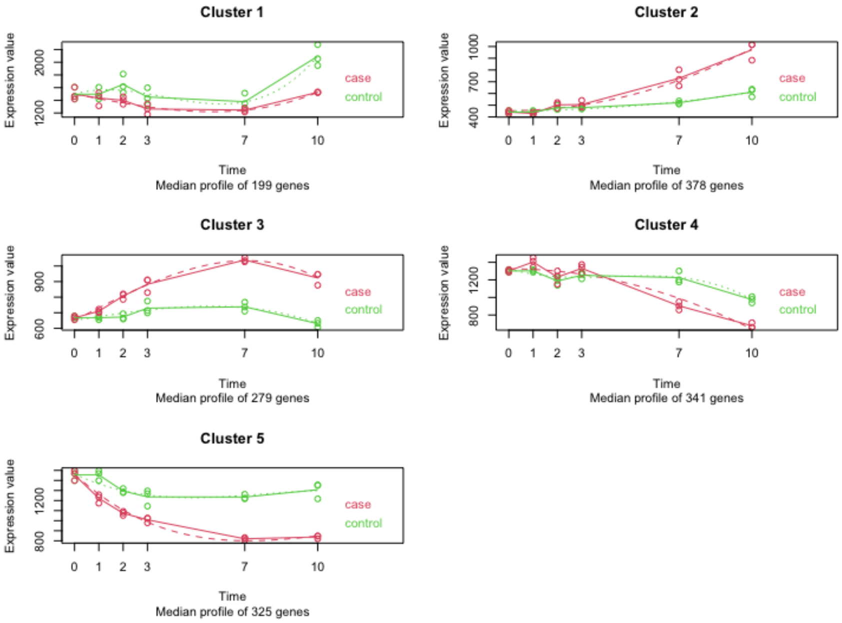


**kcluster=6.**

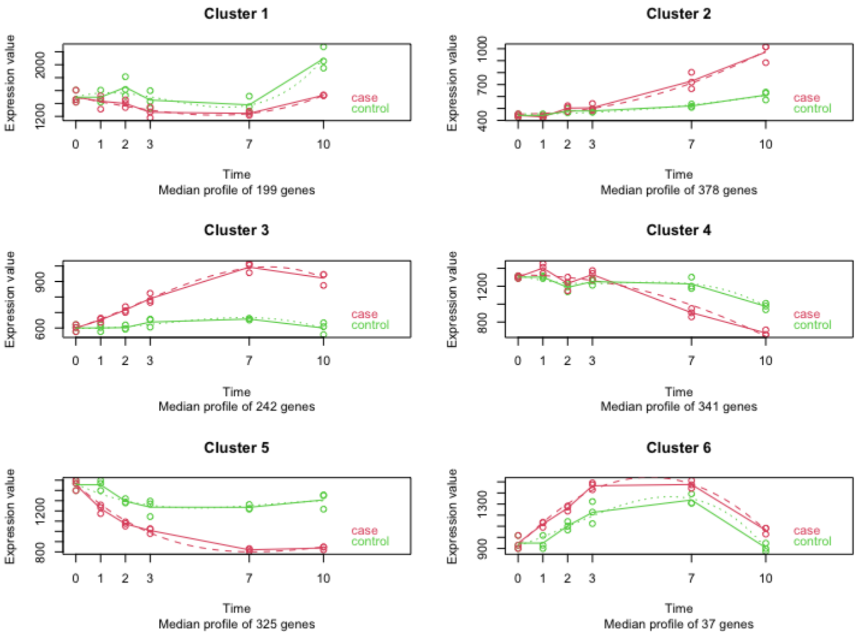

Cluster 1
Expression value
Time
Median profile of 199 genes
case
control
Cluster 2
Median profile of 378 genes
Cluster 3
Median profile of 242 genes
Cluster 4
Median profile of 341 genes
Cluster 5
Median profile of 325 genes
Cluster 6
Median profile of 37 genes


**kcluster=7.**

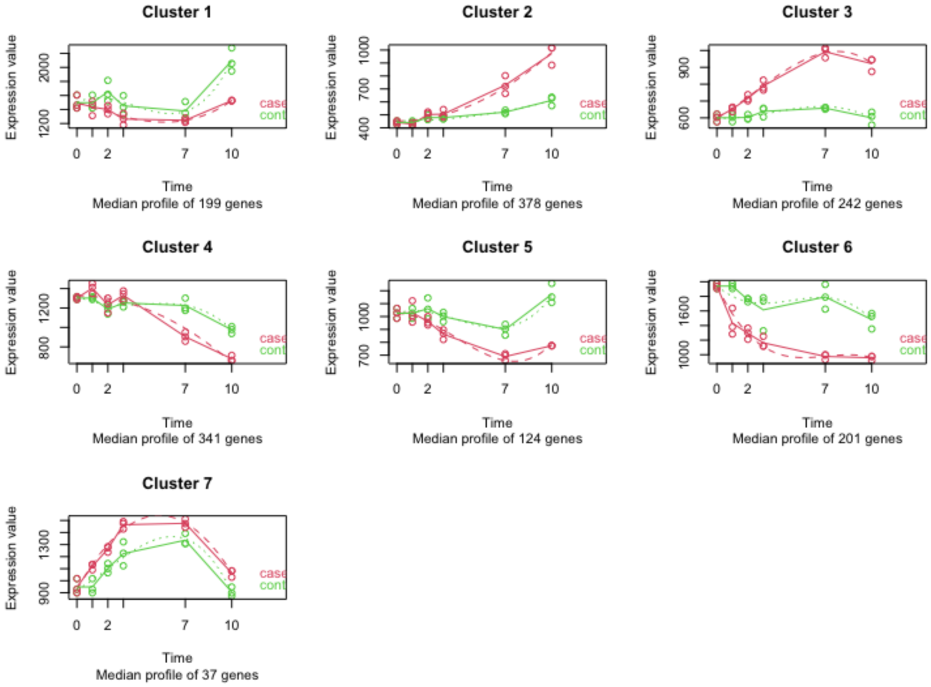

Cluster 1
Expression value
Time
Median profile of 199 genes
Cluster 2
Median profile of 378 genes
Cluster 3
Median profile of 242 genes
Cluster 4
Median profile of 341 genes
Cluster 5
Median profile of 124 genes
Cluster 6
Median profile of 201 genes
Cluster 7
Median profile of 37 genes

**kcluster=8.**

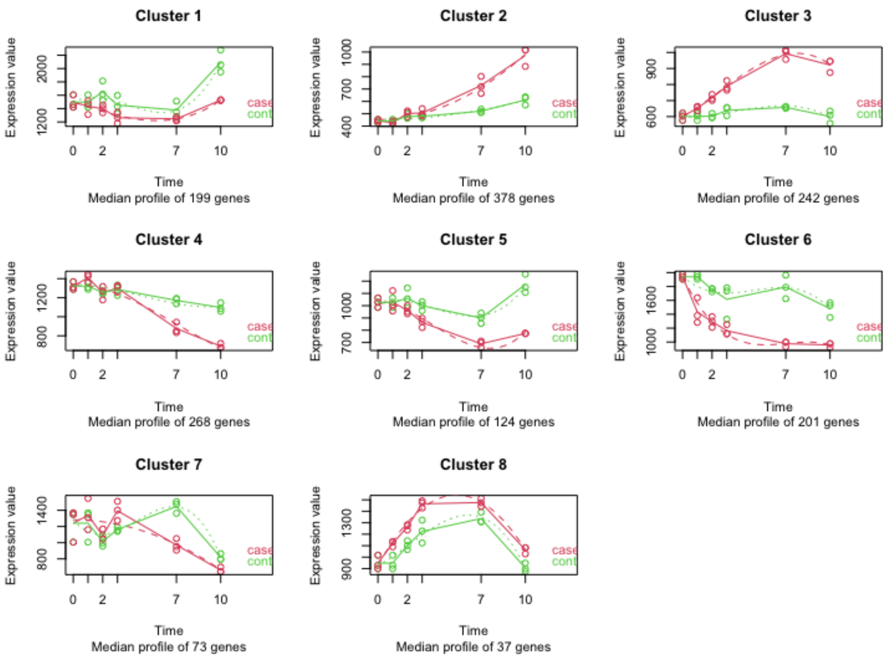


**kcluster=9.**

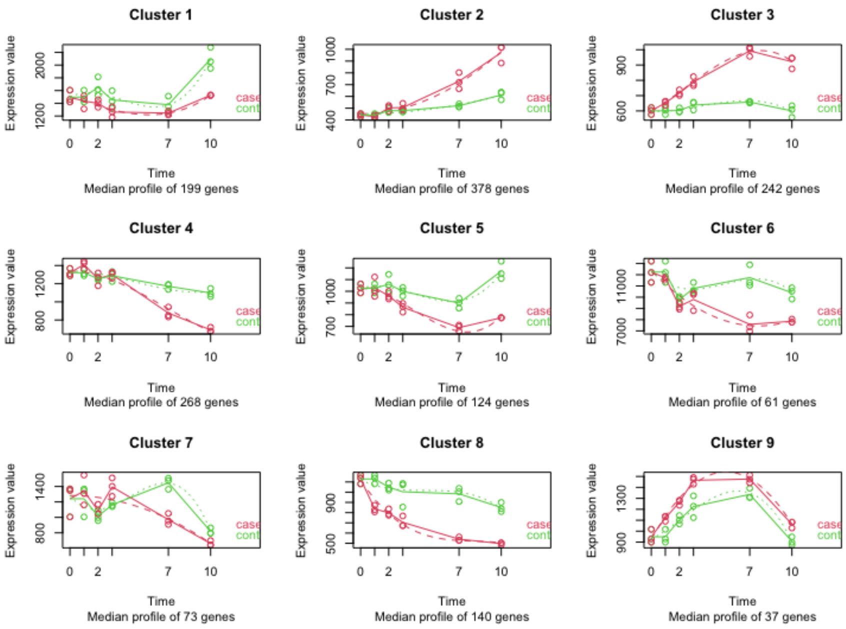

Fig. S2 **Expression clusters** (**maSigpro analysis) for *B. pusilla***

**kcluster=2.**

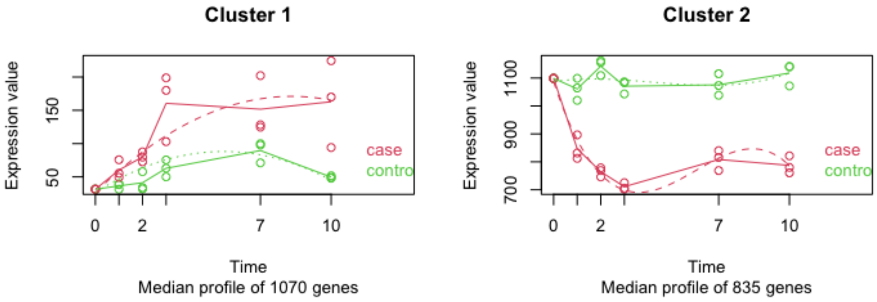


**kcluster=3**

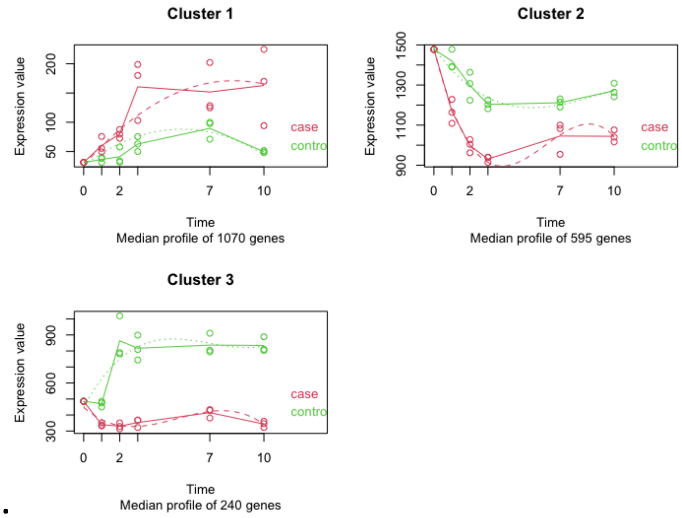


•

**kcluster=4.**

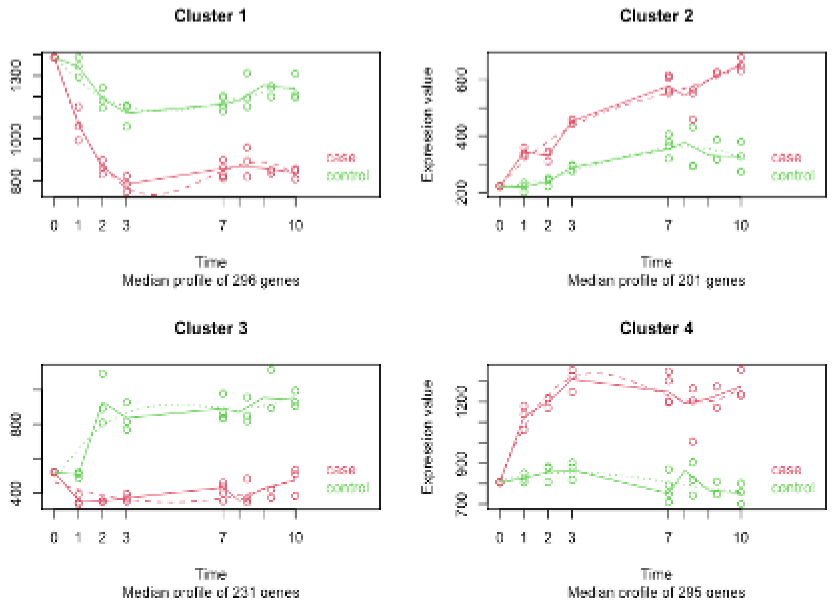


**kcluster=5.**

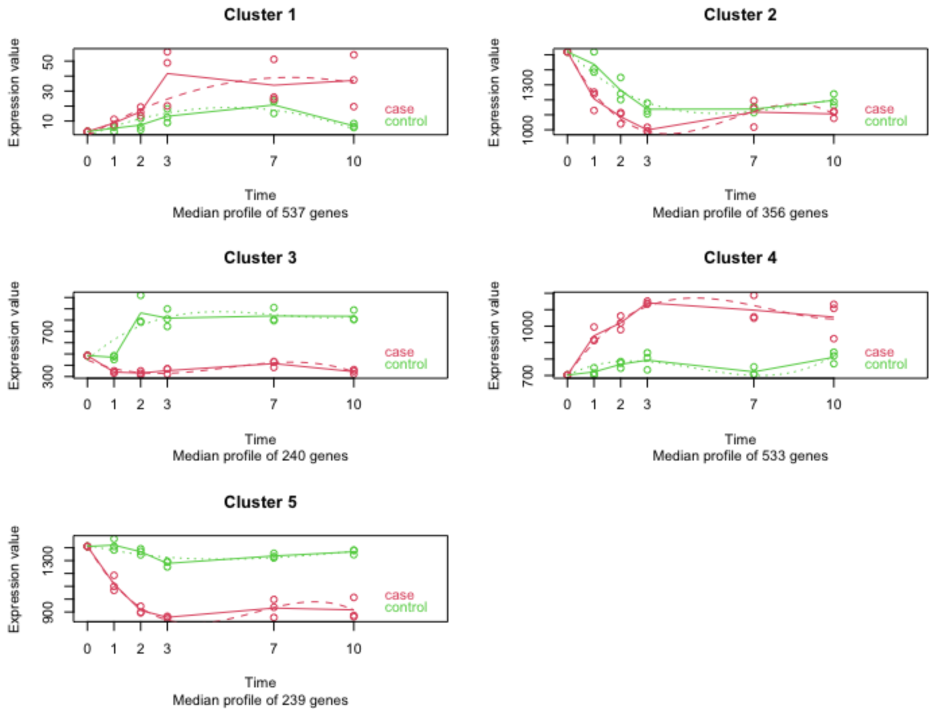


**kcluster=6.**

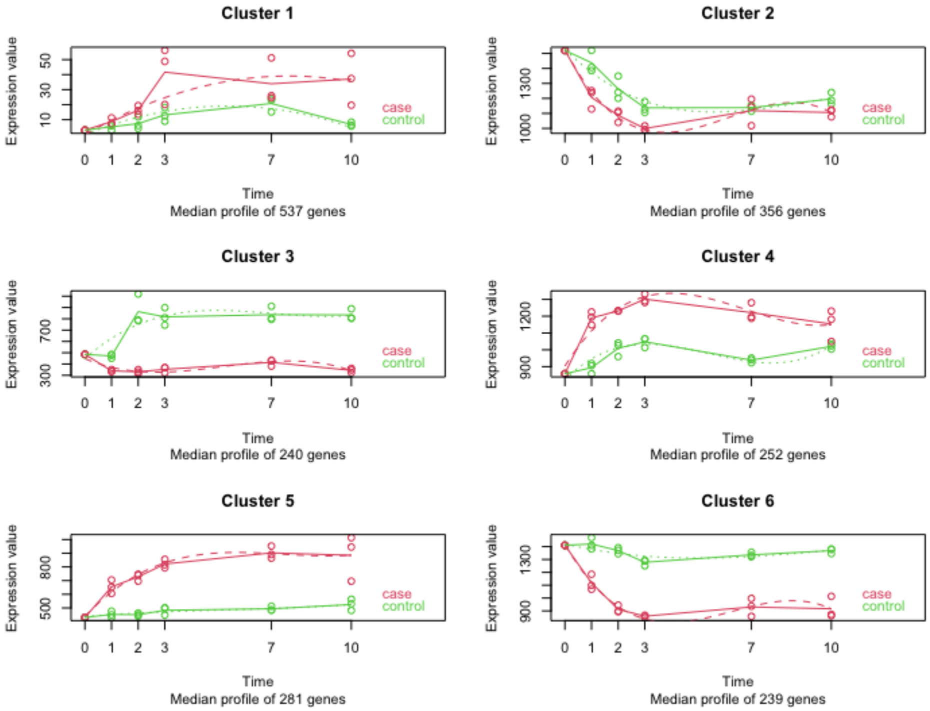
Cluster 1
Expression value
Time
Median profile of 537 genes
case
control
Cluster 2
Median profile of 356 genes
Cluster 3
Median profile of 240 genes
Cluster 4
Median profile of 252 genes
Cluster 5
Median profile of 281 genes
Cluster 6
Median profile of 239 genes

**kcluster=7.**

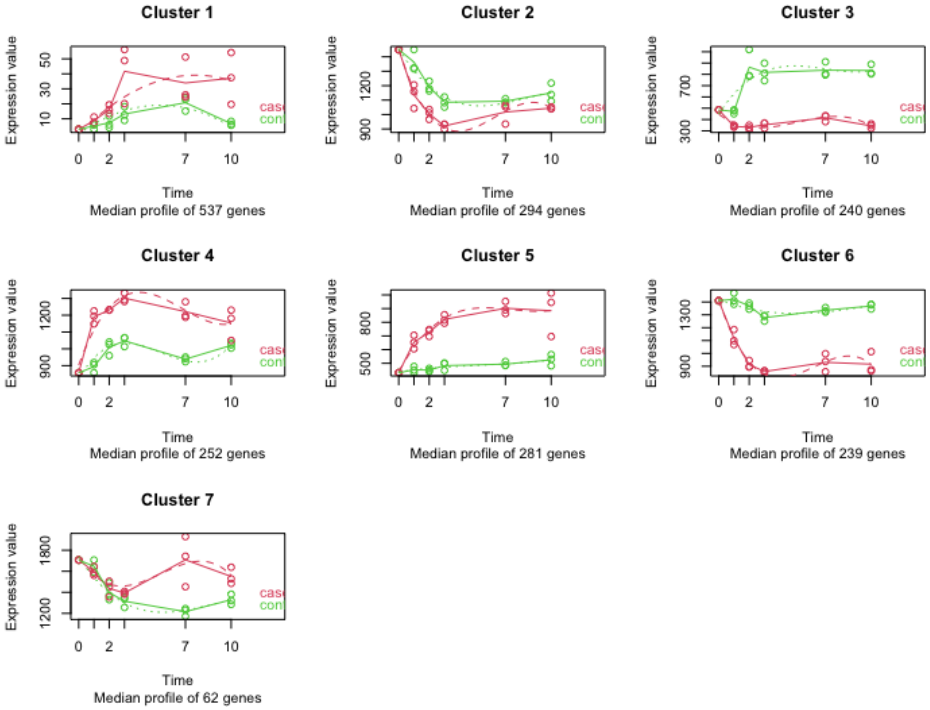
Cluster 1
Expression value
Time
Median profile of 537 genes
Cluster 2
Median profile of 294 genes
Cluster 3
Median profile of 240 genes
Cluster 4
Median profile of 252 genes
Cluster 5
Median profile of 281 genes
Cluster 6
Median profile of 239 genes
Cluster 7
Median profile of 62 genes

**kcluster=8.**

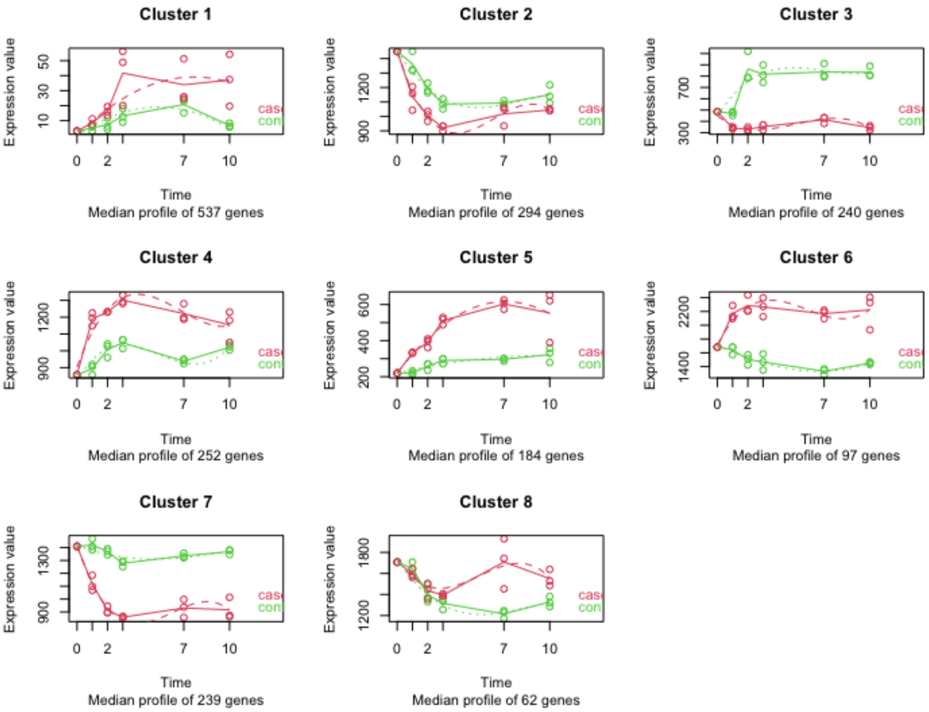


**kcluster=9.**

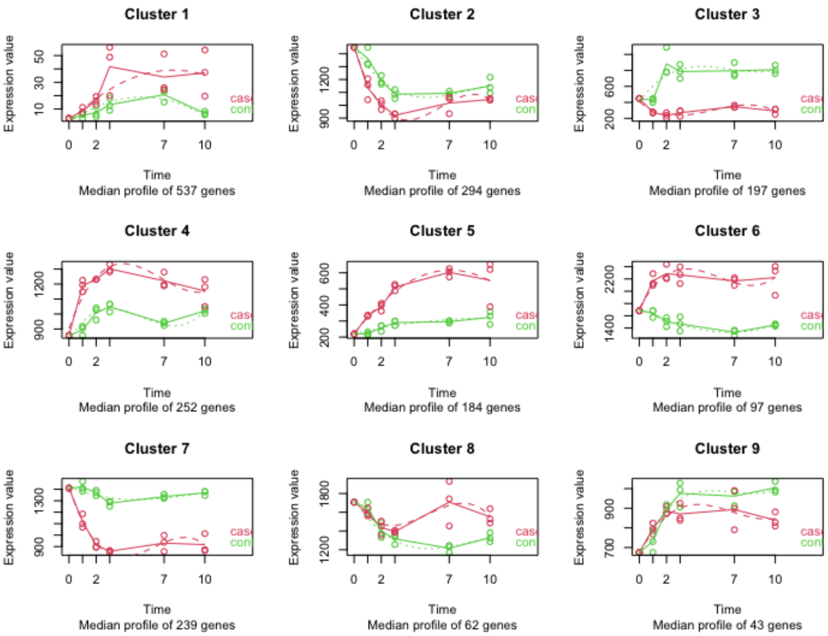

Fig. S3 **Functional enrichment analyses of cluster 1 under nitrogen starvation in *A. agrestis.***

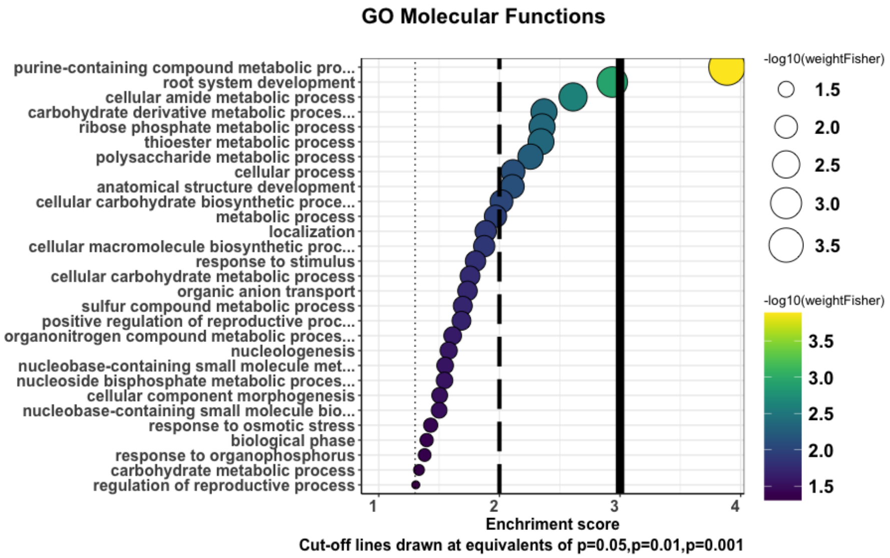


Fig. S4 **Functional enrichment analyses of cluster 2 under nitrogen starvation in *A. agrestis.*.**

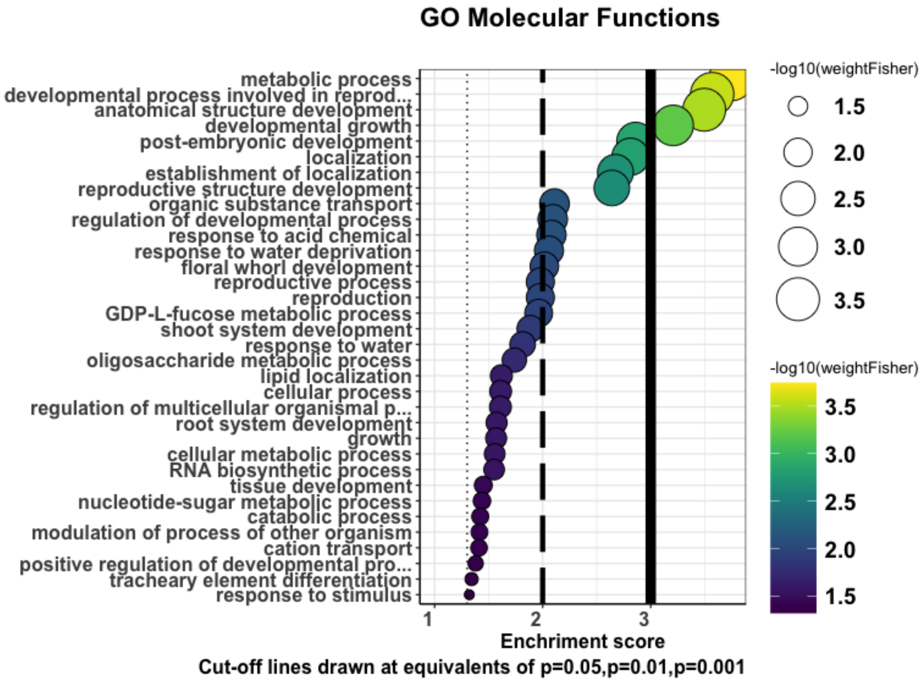

Fig. S5 **Functional enrichment analyses of cluster 3 under nitrogen starvation in *A. agrestis.*.**

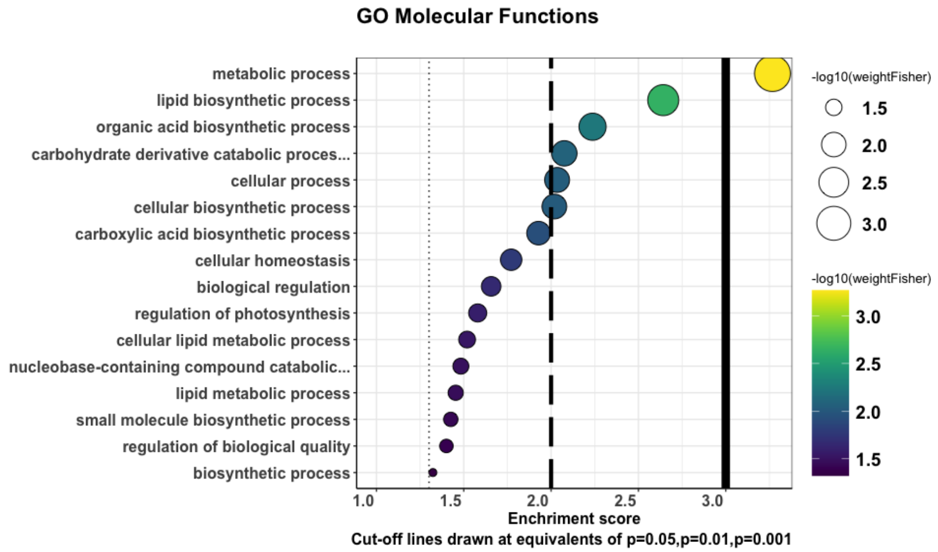


Fig. S6 **Functional enrichment analyses of cluster 4 under nitrogen starvation in *A. agrestis.*.**

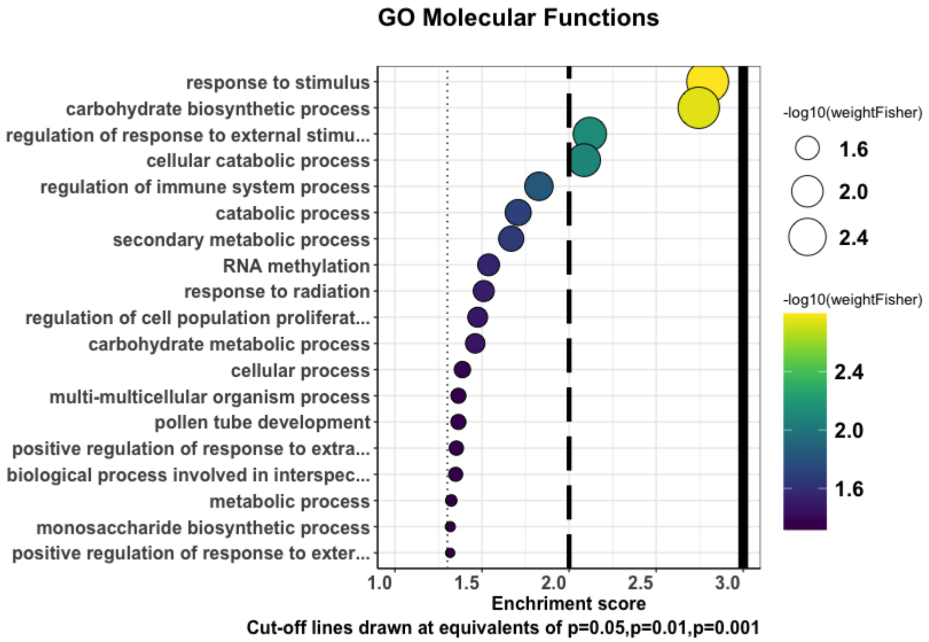

Fig. S7 **Functional enrichment analyses of cluster 1 under nitrogen starvation in *B. pusilla.***

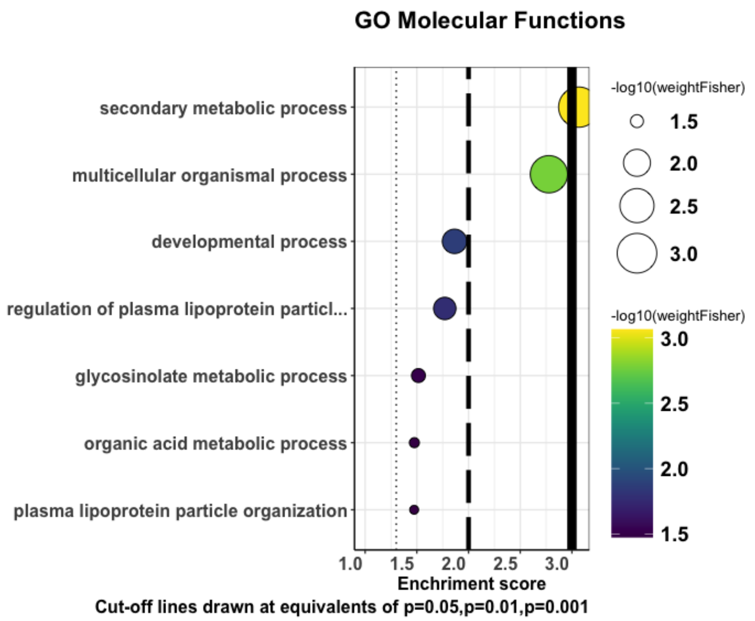


Fig. S8 **Functional enrichment analyses of cluster 2 under nitrogen starvation in *B. pusilla***.

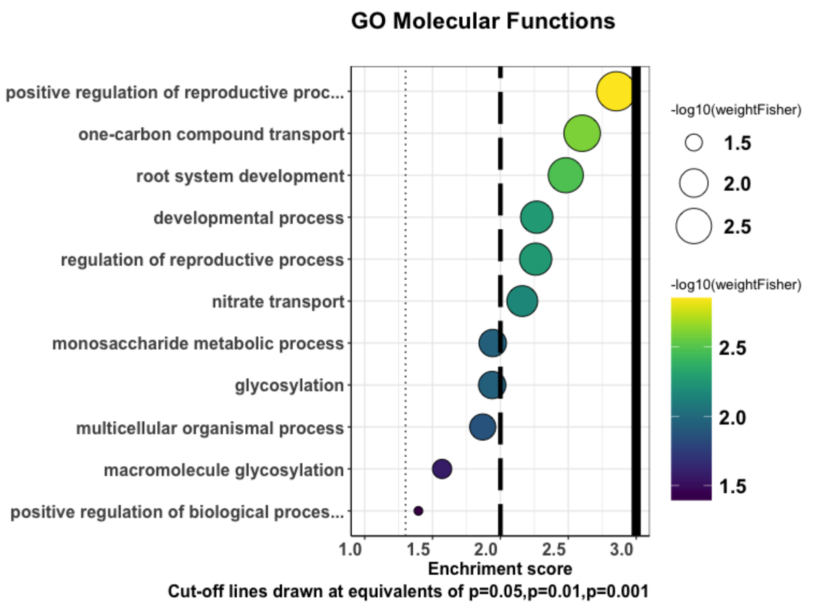


Fig. S9 **Functional enrichment analyses of cluster 3 under nitrogen starvation in *B. pusilla***.

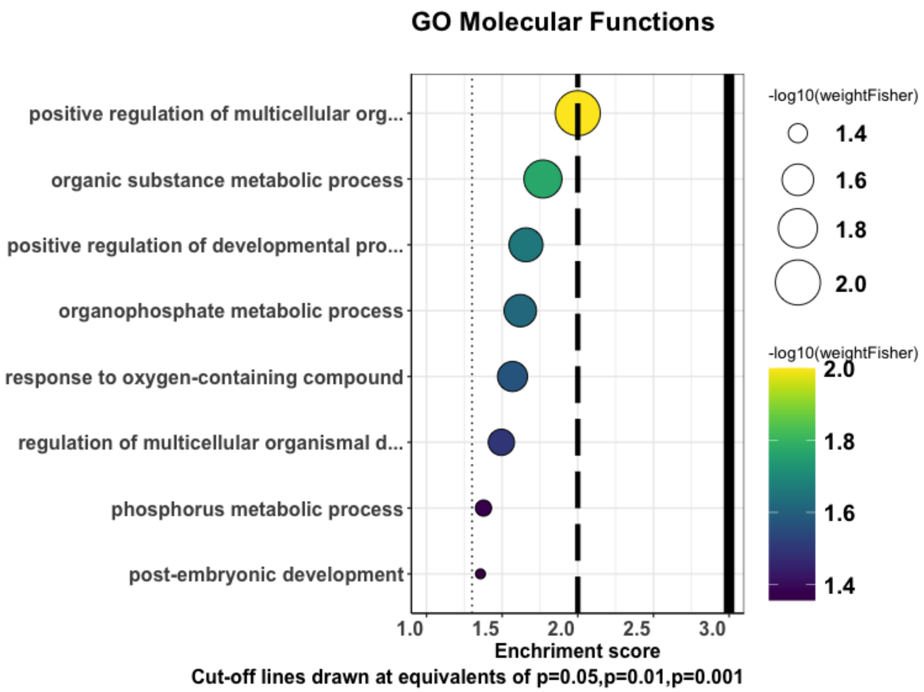


Fig. S10 **Functional enrichment analyses of cluster 4 under nitrogen starvation in *B. pusilla***.

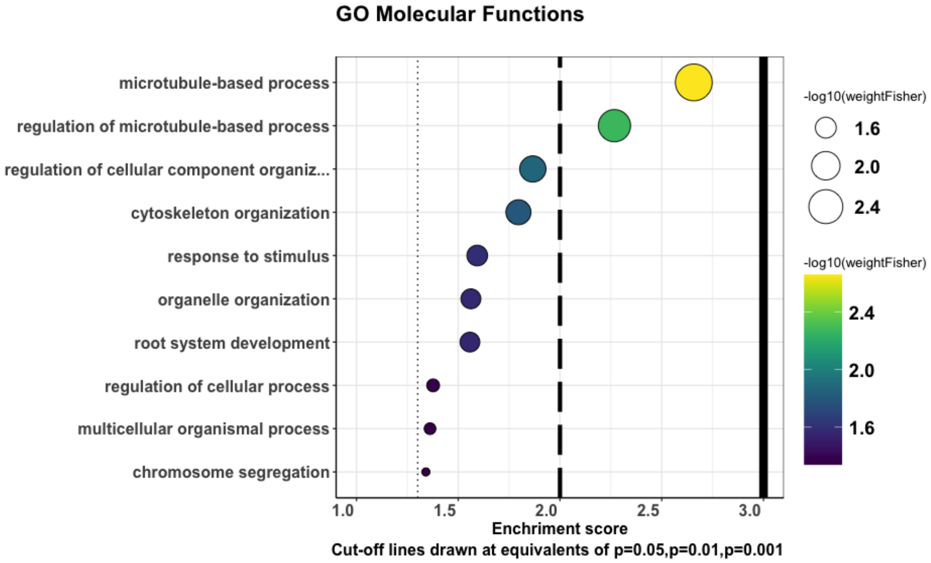
