## Supplementary material for "Nitrogen starvation response in hornworts and liverworts provides little evidence for complex priming to the cyanobiont": Draft_genome

### *Supplementary File 1:* Draft genome assembly of *Blasia pusilla*

**Article title:** Nitrogen starvation response in hornworts and liverworts provides little evidence for complex priming to the cyanobiont

**Authors:** *Yuling Yue, Gaurav Sablok*, *Anna Neubauer, Jaakko Hyvönen*, *Péter Szövényi*

BUSCO scores for all the species we compared (Kirbis et al., 2020)

| | Complete | | | Fragmented | Missing |
|---|---|---|---|---|---|
| | Total | Single-copy | Duplicated | | |
| **Embryophyta (*n* = 1614)** | | | | | |
| *F. hygrometrica* | 1300 (80.5%) | 1101 (68.2%) | 199 (12.3%) | 43 (2.7%) | 271 (16.8%) |
| *M. polymorpha* | 1412 (87.5%) | 1385 (85.8%) | 27 (1.7%) | 20 (1.2%) | 182 (11.3%) |
| *P. patens* | 1423 (88.1%) | 1174 (72.7%) | 249 (15.4%) | 28 (1.7%) | 163 (10.2%) |
| *P. schreberi* | 736 (45.6%) | 540 (33.5%) | 196 (12.1%) | 278 (17.2%) | 600 (37.2%) |
| *S. fallax* | 1447 (89.7%) | 1257 (77.9%) | 190 (11.8%) | 22 (1.4%) | 145 (8.9%) |
| **Viridiplantae (*n* = 425)** | | | | | |
| *F. hygrometrica* | 390 (91.8%) | 336 (79.1%) | 54 (12.7%) | 7 (1.6%) | 28 (6.6%) |
| *M. polymorpha* | 411 (96.7%) | 409 (96.2%) | 2 (0.5%) | 3 (0.7%) | 11 (2.6%) |
| *P. patens* | 416 (97.8%) | 361 (84.9%) | 55 (12.9%) | 1 (0.2%) | 8 (2.0%) |
| *P. schreberi* | 227 (53.4%) | 160 (37.6%) | 67 (15.8%) | 107 (25.2%) | 91 (21.4%) |
| *S. fallax* | 417 (98.1%) | 375 (88.2%) | 42 (9.9%) | 2 (0.5%) | 6 (1.4%) |

*For comparison, BUSCO statistics for the Marchantia polymorpha, Physcomitrium patens, Pleurozium schreberi, and Sphagnum fallax genomes are shown. Data was retrieved from Phytozome v12 (Goodstein et al., 2012) and analyzed using the Viridiplantae and Embryophyta BUSCO datasets obtained from OrthoDB v10.1.*

BUSCO scores for the blasia draft genome assembly before blobtools fitering and after bolbtools fitering

Genome: blasia_2020_hybrid_polishing.fasta

The lineage dataset is: viridiplantae_odb10

***** Results: *****

C:86.9%[S:79.8%,D:7.1%],F:4.0%,M:9.1%,n:425

369 Complete BUSCOs (C)

339 Complete and single-copy BUSCOs (S)

30 Complete and duplicated BUSCOs (D)

17 Fragmented BUSCOs (F)

39 Missing BUSCOs (M)

425 Total BUSCO groups searched

The lineage dataset is: embryophyta_odb10

***** Results: *****

C:75.3%[S:71.6%,D:3.7%],F:5.0%,M:19.7%,n:1614

1216 Complete BUSCOs (C)

1156 Complete and single-copy BUSCOs (S)

60 Complete and duplicated BUSCOs (D)

81 Fragmented BUSCOs (F)

317 Missing BUSCOs (M)

1614 Total BUSCO groups searched

Genome: blasia_2020_hybrid_polishing_FILTEREDonlyPLANT19Jan2022.fasta (after blobtools filtering)

The lineage dataset is: viridiplantae_odb10

***** Results: *****

C:79.7%[S:79.5%,D:0.2%],F:4.7%,M:15.6%,n:425

339 Complete BUSCOs (C)

338 Complete and single-copy BUSCOs (S)

1 Complete and duplicated BUSCOs (D)

20 Fragmented BUSCOs (F)

66 Missing BUSCOs (M)

425 Total BUSCO groups searched

The lineage dataset is: embryophyta_odb10

***** Results: *****

C:71.0%[S:69.6%,D:1.4%],F:4.5%,M:24.5%,n:1614

1146 Complete BUSCOs (C)

1123 Complete and single-copy BUSCOs (S)

23 Complete and duplicated BUSCOs (D)

72 Fragmented BUSCOs (F)

396 Missing BUSCOs (M)

1614 Total BUSCO groups searched

#### Quast results:

Genome: blasia_2020_hybrid_polishing.fasta

| **Statistics without reference** | **blasia_2020_hybrid_polishing** |
|---|---|
| # contigs | 3569 |
| # contigs (>= 0 bp) | 3569 |
| # contigs (>= 1000 bp) | 3037 |
| # contigs (>= 5000 bp) | 2714 |
| # contigs (>= 10000 bp) | 2562 |
| # contigs (>= 25000 bp) | 2353 |

| | |
|---|---|
| # contigs (>= 50000 bp) | 1951 |
| Largest contig | 6594954 |
| Total length | 607953238 |
| Total length (>= 0 bp) | 607953238 |
| Total length (>= 1000 bp) | 607601461 |
| Total length (>= 5000 bp) | 606728275 |
| Total length (>= 10000 bp) | 605631448 |
| Total length (>= 25000 bp) | 601920055 |
| Total length (>= 50000 bp) | 586818066 |
| N50 | 486748 |
| N90 | 94585 |
| auN | 950925 |
| L50 | 302 |
| L90 | 1389 |
| GC (%) | 49.68 |
| **Mismatches** | |
| # N's per 100 kbp | 0.09 |
| # N's | 546 |

Genome: blasia_2020_hybrid_polishing_FILTEREDonlyPLANT19Jan2022.fasta (after blobtools filtering)

| **Statistics without reference** | **blasia_2020_hybrid_polishing_...** |
|---|---|
| # contigs | 1456 |
| # contigs (>= 0 bp) | 1456 |
| # contigs (>= 1000 bp) | 1403 |
| # contigs (>= 5000 bp) | 1363 |
| # contigs (>= 10000 bp) | 1330 |
| # contigs (>= 25000 bp) | 1251 |
| # contigs (>= 50000 bp) | 1105 |
| Largest contig | 2605490 |
| Total length | 347238166 |
| Total length (>= 0 bp) | 347238166 |
| Total length (>= 1000 bp) | 347204272 |
| Total length (>= 5000 bp) | 347092030 |
| Total length (>= 10000 bp) | 346842215 |
| Total length (>= 25000 bp) | 345433237 |
| Total length (>= 50000 bp) | 340065909 |

| | |
|---|---|
| N50 | 452662 |
| N90 | 125550 |
| auN | 571767 |
| L50 | 224 |
| L90 | 774 |
| GC (%) | 44.12 |
| **Mismatches** | |
| # N's per 100 kbp | 0.16 |
| # N's | 546 |
