## Supplementary material for "Nitrogen starvation response in hornworts and liverworts provides little evidence for complex priming to the cyanobiont": Transcriptomic_reference

### *Supplementary File 2: Evaluation of the Blasia transcriptome reference*

**Article title:** Nitrogen starvation response in hornworts and liverworts provides little evidence for complex priming to the cyanobiont

**Authors:** *Yuling Yue, Gaurav Sablok*, *Anna Neubauer, Jaakko Hyvönen*, *Péter Szövényi*

–

Denovo assembled transcriptome

RNA_seq_Blasia_trinity_denovo_with_v2.13.2_kmecovuntouched.Trinity.fasta

```
################################
## Counts of transcripts, etc.
################################
Total trinity 'genes':  181502
Total trinity transcripts:      290809
Percent GC: 48.36

########################################
 Stats based on ALL transcript contigs:
 ########################################

Contig N10: 7876
Contig N20: 6223
Contig N30: 5145
Contig N40: 4304
Contig N50: 3568

Median contig length: 652
Average contig: 1608.99
Total assembled bases: 467910189


#####################################################
## Stats based on ONLY LONGEST ISOFORM per 'GENE':
 #####################################################

Contig N10: 7019
Contig N20: 5271
Contig N30: 4106
Contig N40: 3173
Contig N50: 2362

Median contig length: 356
Average contig: 940.65
Total assembled bases: 170730752
```

at least one complete CDS sequence : All_transcripts_withCOMPLETECDS_FINALOK.fasta

Transcriptome Contig Nx and ExN50 stats

Error, cannot decipher gene identifier from acc: asmbl_1

################################

#### Counts of transcripts, etc.

################################

Total trinity 'genes':    180418

Total trinity transcripts:          180418

Percent GC: 47.01

########################################

Stats based on ALL transcript contigs:

########################################

Contig N10: 10450

Contig N20: 8462

Contig N30: 7305

Contig N40: 6428

Contig N50: 5681

Median contig length: 4086

Average contig: 4544.05

Total assembled bases: 819827674

- note: not reporting gene-based longest isoform info since couldn't parse Trinity accession info.

read alignment statistics for paired-end reads:

20538334 reads; of these:
  20538334 (100.00%) were paired; of these:

```
1957048 (9.53%) aligned concordantly 0 times
751122 (3.66%) aligned concordantly exactly 1 time
17830164 (86.81%) aligned concordantly >1 times
 ----
 1957048 pairs aligned concordantly 0 times; of these:
 24231 (1.24%) aligned discordantly 1 time
----
 1932817 pairs aligned 0 times concordantly or discordantly; of these:
 3865634 mates make up the pairs; of these:
 2066006 (53.45%) aligned 0 times
 54136 (1.40%) aligned exactly 1 time
1745492 (45.15%) aligned >1 times
 94.97% overall alignment rate
```

Busco scores:

#INFO:

```
--------------------------------------------------
|Results from dataset viridiplantae_odb10          |
--------------------------------------------------
|C:99.8%[S:2.6%,D:97.2%],F:0.2%,M:0.0%,n:425 |
|424    Complete BUSCOs (C)                        |
|11     Complete and single-copy BUSCOs (S)        |
|413    Complete and duplicated BUSCOs (D)         |
|1      Fragmented BUSCOs (F)                      |
|0      Missing BUSCOs (M)                         |
|425    Total BUSCO groups searched                |
--------------------------------------------------
```

#INFO:

```
--------------------------------------------------
|Results from dataset embryophyta_odb10            |
--------------------------------------------------
|C:87.1%[S:1.7%,D:85.4%],F:2.5%,M:10.4%,n:1614     |
|1406   Complete BUSCOs (C)                        |
|28     Complete and single-copy BUSCOs (S)        |
|1378   Complete and duplicated BUSCOs (D)         |
|40     Fragmented BUSCOs (F)                      |
```

| 168 Missing BUSCOs (M) |
| 1614 Total BUSCO groups searched |
--------------------------------------------------

All_transcripts_withCOMPLETECDS_FINALOK_cleangenes.fasta

(after removing yeast Viruses, Bacteria, Archeae, Opisthokonta and other Eukaryota)

Error, cannot decipher gene identifier from acc: asmbl_1

################################

#### Counts of transcripts, etc.

################################

Total trinity 'genes':    156239

Total trinity transcripts:          156239

Percent GC: 46.76

########################################

Stats based on ALL transcript contigs:

########################################

Contig N10: 10570

Contig N20: 8589

Contig N30: 7426

Contig N40: 6549

Contig N50: 5816

Median contig length: 4393

Average contig: 4858.91

Total assembled bases: 759151709

- note: not reporting gene-based longest isoform info since couldn't parse Trinity accession info.

Busco scores:

```
--------------------------------------------------
	|Results from dataset viridiplantae_odb10	 |
	--------------------------------------------------
	|C:97.7%[S:2.4%,D:95.3%],F:0.2%,M:2.1%,n:425	  |
	|415	Complete BUSCOs (C)			  |
	|10	Complete and single-copy BUSCOs (S)	  |
	|405	Complete and duplicated BUSCOs (D)	  |
	|1	Fragmented BUSCOs (F)			  |
	|9	Missing BUSCOs (M)			  |
	|425	Total BUSCO groups searched		  |
	--------------------------------------------------
--------------------------------------------------
	|Results from dataset embryophyta_odb10	  |
	--------------------------------------------------
	|C:85.1%[S:1.9%,D:83.2%],F:2.7%,M:12.2%,n:1614	  |
	|1373	Complete BUSCOs (C)			  |
	|30	Complete and single-copy BUSCOs (S)	  |
	|1343	Complete and duplicated BUSCOs (D)	  |
	|43	Fragmented BUSCOs (F)			  |
	|198	Missing BUSCOs (M)			  |
	|1614	Total BUSCO groups searched		  |
	--------------------------------------------------
```
